# Deprivation-induced plasticity in the early central circuits of the rodent visual, auditory, and olfactory systems: a systematic review and meta-analysis of the literature

**DOI:** 10.1101/2023.09.04.556170

**Authors:** Li Huang, Francesca Hardyman, Megan Edwards, Elisa Galliano

## Abstract

Activity-dependent neuronal plasticity is crucial for animals to adapt to dynamic sensory environments. Traditionally, research on activity dependent-plasticity has used sensory deprivation approaches in animal models, and it has focused on its effects in primary sensory cortices. However, emerging evidence emphasizes the importance of activity-dependent plasticity both in the sensory organs and in sub-cortical regions where cranial nerves relay information to the brain. Additionally, a critical question arises: do different sensory modalities share common cellular mechanisms for deprivation-induced plasticity at these central entry-points? Furthermore, does the duration of deprivation correlate with specific plasticity mechanisms? This study aims to systematically review and meta-analyse research papers that investigated visual, auditory, or olfactory deprivation in rodents. Specifically, it explores the consequences of sensory deprivation in homologous regions at the first central synapse after the cranial nerve: vision—lateral geniculate nucleus and superior colliculus; audition— ventral and dorsal cochlear nucleus; olfaction—olfactory bulb. The systematic search yielded 91 research papers (39 vision, 22 audition, 30 olfaction), revealing significant heterogeneity in publication trends, experimental methods of inducing deprivation, measures of deprivation-induced plasticity, and reporting, across the three sensory modalities. Nevertheless, despite these methodological differences, commonalities emerged when correlating the plasticity mechanisms with the duration of the sensory deprivation. Following short-term deprivations (up to 1 day) all three systems showed reduced activity levels and increased disinhibition. Medium-term deprivation (1 day to a week) induced greater glial involvement and synaptic remodelling. Long-term deprivation (over a week) predominantly led to macroscopic structural changes including tissue shrinkage and apoptosis. These findings underscore the importance of standardizing methodologies and reporting practices. Additionally, they highlight the value of cross-modals synthesis for understanding how the nervous system, including peripheral, pre-cortical, and cortical areas, respond to and compensate for sensory inputs loss.

## Introduction

Animals rely on sophisticated sensory organs to effectively perceive and interact with their surroundings. These sensory organs can convert various environmental stimuli, such as electromagnetic waves, mechanical pressure, and chemicals, into trains of action potentials that are relayed and computed in dedicated brain areas. The disruption of the sensory transduction cascade is a common occurrence attributable to factors such as trauma, ischemia, viral infection, and aging(1–6). If left unattended, sudden sensory loss can significantly impact an individual’s behaviour and wellbeing. Consequently, the nervous system must promptly adopt strategies to compensate for such losses. Unlike organs like bones or skin, the adult brain cannot regenerate damaged peripheral sensors or central neurons, with only the olfactory system being a notable exception(7,8). However, neurons can partially counteract the loss of sensory information by engaging a range of activity-dependent plasticity mechanisms(9). These encompass both functional and structural changes at synapses, as well as adjustments in the intrinsic excitability and firing rates of neurons(10–12). Furthermore, glial cells also play a pivotal role in facilitating neuronal plasticity(13,14). The investigation of the mechanisms behind deprivation-induced adaptive plasticity across different timeframes, from immediate sensory loss to subsequent functional recovery, not only enhances our fundamental understanding of how neural circuits adapt to changing sensory inputs but also holds great significance for translational research in improving recovery after sudden sensory loss (15,16).

Since the seminal experiments of Hubel and Wiesel in monocularly deprived kittens(17), multiple studies have dissected the mechanisms of deprivation-induced plasticity in animal models. This extensive body of work has largely examined the deprivation effects in primary sensory cortices during developmental critical periods and adulthood(18,19). While less emphasis has been placed on pre-cortical regions(20,21), it’s important to note that changes in these areas have a cascading impact on cortical adaptation and processing. Furthermore, existing studies in both cortex and pre-cortical areas have primarily focused on individual sensory modalities, making meaningful cross-modal comparisons challenging due to substantial experimental variability. This variability arises from factors such as the animals’ developmental stage, the experimental model, and heterogeneous methods for inducing adaptive plasticity through sensory deprivation(22,23). Additionally, diversity in experimental design and result reporting complicates efforts to establish overarching principles governing the recruitment of different plasticity mechanisms in excitatory and inhibitory neurons, as well as glial cells. Synthesizing general principles is further hindered by anatomical and physiological diversity in the various sensory pathways, including differences in sensory organ complexity, transduction mechanisms, and the number of pre-cortical relays. To overcome these complications, this study focuses on deprivation-induced plasticity in anatomically homologous subcortical hubs in the olfactory, visual and auditory pathways. The olfactory bulb (OB), lateral geniculate nucleus and superior colliculus (LGN and SC), and the dorsal and ventral cochlear nuclei (DCN and VCN) receive the primary synapse in the brain made by the respective cranial nerves (Figure 1). While their circuit architectures differs granularly, these five circuits share multiple features(20,24–29). Principal neurons receive glutamatergic inputs from the olfactory, optic, or cochlear nerves, and send their axons to higher processing areas (from which they receive feedback projections which are beyond the scope of this study). Importantly, all five circuits heavily feature local inhibitory interneurons modulating information transfer, with additional excitatory interneurons described in both OB and DCN.

**Figure 1.**
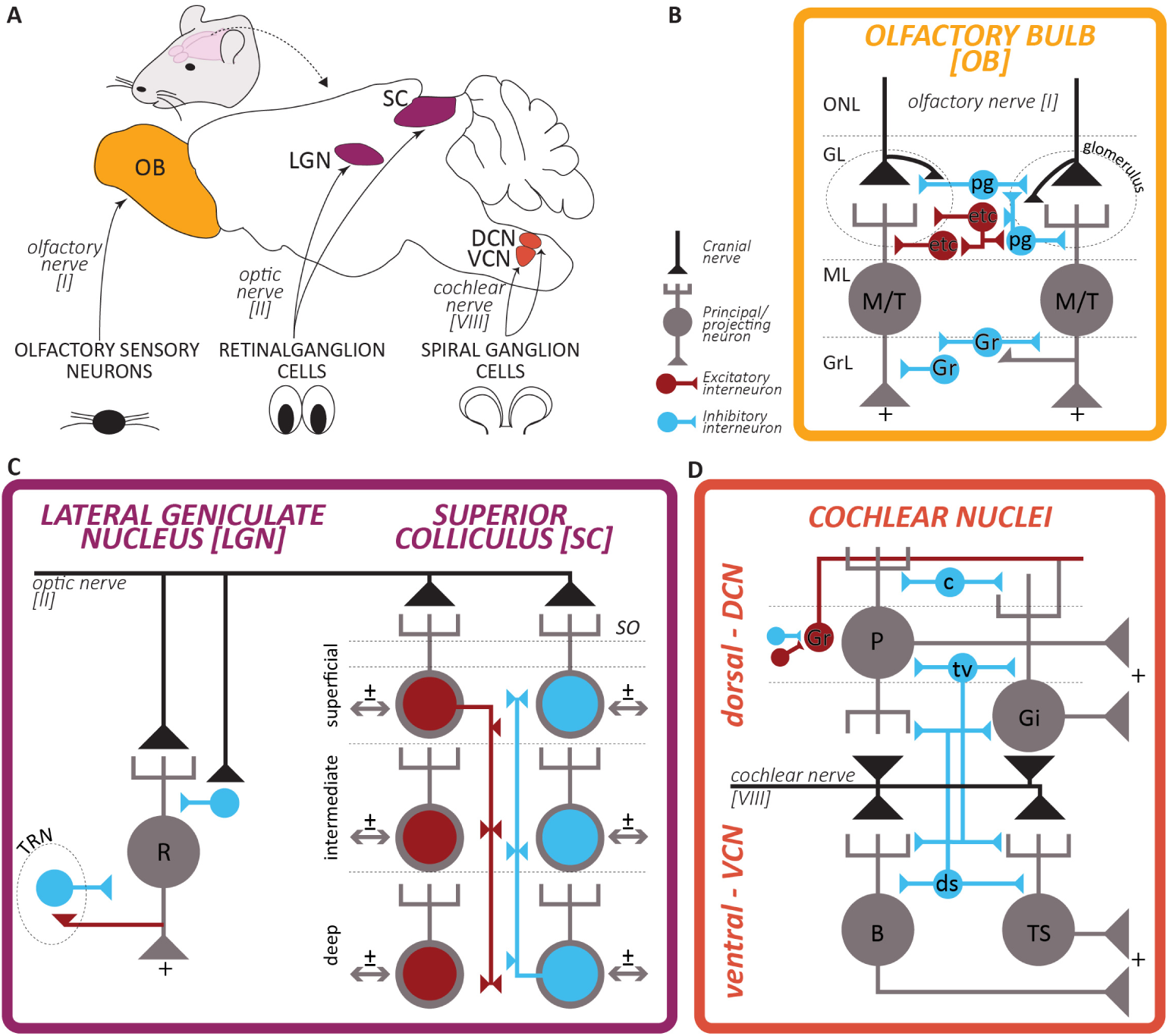
Architecture of the early olfactory, visual, and auditory pathways. (A) Schematic representation of the mouse brain and location of the olfactory bulb (OB), lateral geniculate nucleus (LGN; dorsal, dLGN, and ventral, vLGN, combined), superior colliculus (SC), dorsal and ventral cochlear nuclei (DCN and VCN), and their respective cranial nerve inputs from the sensory organs. **(B-D)** Simplified circuitry of the early central circuits processing olfactory, visual, and auditory information summarized and adapted from (20,24–29). Black line and triangle indicate the cranial nerve endings, grey cells are principal neurons projecting outside these early circuits to higher processing areas, red and blue cells are local interneurons, respectively excitatory and inhibitory. For ease of representation, the many central inputs to these circuits are not depicted. In the bulbar circuit: ONL = olfactory nerve layer; GL= glomerular layer; ML = mitral layer; GrC = granule cells layer; M/T = mitral/tufted cell; pg = periglomerular cell; etc = external tufted cell; gr = granule cell. In the geniculate circuit: R = relay neuron; TRN = thalamic reticular nucleus. In the collicular circuit: SO = stratum opticum; note that the exact circuitry has not been fully resolved, and that all layers send projections outside the SC. In the cochlear nuclei circuit: P = pyramidal (or fusiform); Gi = giant cell; B =bushy cell; TS = t-stellate cell; ds = d-stellate cell; tv = tubercoloventral (or vertical) cell; c = cartwheel cell; Gr = granule cell with its axon called parallel fibre.

**Figure 2.**
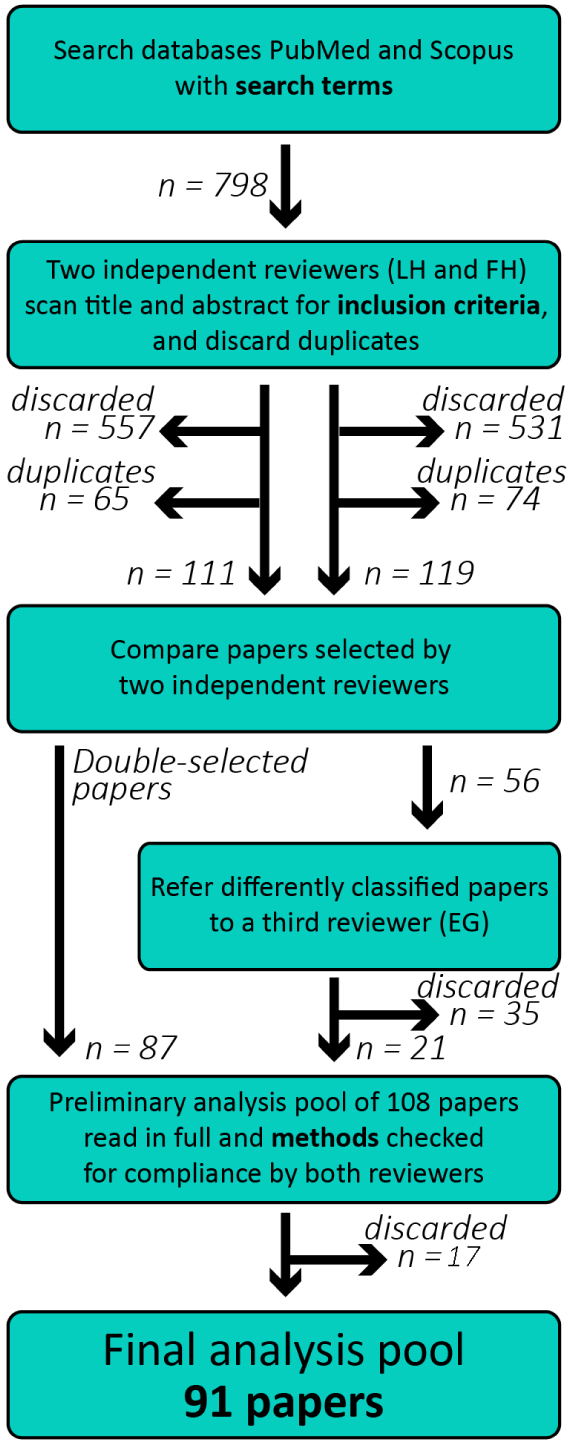
Strategy for literature search and papers selection. Flow chart indicating the number of articles returned by the parallel searches in the databases PubMed and Scopus using the search terms detailed in the text, and their subsequent selection by two independent scrutineers.

To disentangle developmentally-regulated plasticity from activity-dependent plasticity in adulthood, we exclusively considered *in vivo* deprivation studies in post-weaning rodents (21 days or older). Our systematic search across two databases returned 91 articles which employed visual, auditory, or olfactory deprivations spanning durations from 30 minutes to over a year and employing a range of experimental from transcriptomics to behavioural assays. This meta-research study pursued two primary objectives. First, we aimed to elucidate the characteristics of the literature and provide recommendations for designing, executing, and reporting sensory deprivation experimental approaches in rodent models. Second, we sought to identify commonalities in pre-cortical plasticity across the three senses, facilitating the synthesis of generalizable principles. Such insights can inform analogous approaches in other systems (*e.g.* in cortex, following sensory enrichment), as well as translational research on recovery from sudden sensory loss.

## Results

### Literature characteristics: publication trends over time

First, we analysed the publication trends over time across vision, audition, olfaction in the qualifying papers which investigated deprivation-induced plasticity in cranial nerve receiving areas. We found no significant trend in the number of publication over time (Figure 3A, simple linear regression; R2=0.06; F(1,36)=2.34; p=0.14). The mean publications per year was 2.4 papers (SD = 1.7). The number of studies investigating the visual system was higher than in olfaction and audition (cumulative distributions, Figure 3B). The median publication year was calculated for each sense, and vision studies appear to be older than olfaction and audition (Figure 3C, Vision, n=39, median = 2002; Audition, n=22, median = 2009; Olfaction, n=30, median = 2009). In summary, we found that the field of deprivation-induced plasticity is steadily productive, with studies focussing on vision being more numerous.

**Figure 3.**
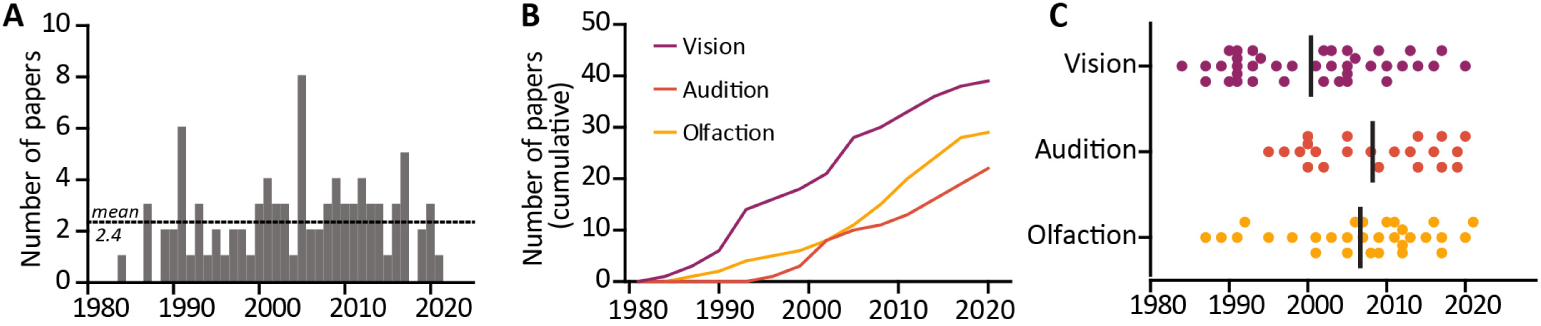
Publication trends over time. (A) No significant trend in the volume of published primary research articles over the 1984-2021 time period. Dotted line represents mean number of publications per year (2.4 mean, 1.7 standard deviation). **(B)** Cumulative distribution of papers published over the 1984-2021 period for each investigated sensory modality. **(C)** Distribution of papers over time for each investigated sensory modality. Dots are individual studies, line indicates median year. Purple = vision, orange = audition, yellow = olfaction.

### Features of the experimental models

Next, we analysed the use of animal models over years and across senses. Most studies used either rats (n=54, of which 35% Wistar and 30% Sprague-Dawley) or mice (n=33, of which 48% C57Bl6 wildtype, 15% wildtype animals in other genetic background, and 30% genetically modified mice). In line with general trends in the field and the advent of numerous commercially available genetically modified lines, the use of mice has significantly increased since the early 2000s (mouse median publication year = 2012; rat median publication year = 2001). Only three studies used other rodent models, namely gerbils (n=1), guinea pigs (n=1) and hamsters (n=2; Fig. 4A). Most studies used exclusively male rodents (41%), and unfortunately many articles did not report the animals’ sex (33%). However, recent years saw an increase of studies using female rodents (Female only, n= 10, median publication year = 2014; Male only: n=37, median publication year =2005; both sexes: n=14, median publication year = 2003; Not reported: n=30, median publication year = 2005; Fig. 4B). Finally, we found no significant differences in the proportion of studies using both sexes among the three sensory modalities, with similar proportions of papers using both sexes (vision=13%, audition=18%, olfaction=17%, chi-squared test X2(2)=1.04; p=0.59; Fig. 4C).

**Figure 4.**
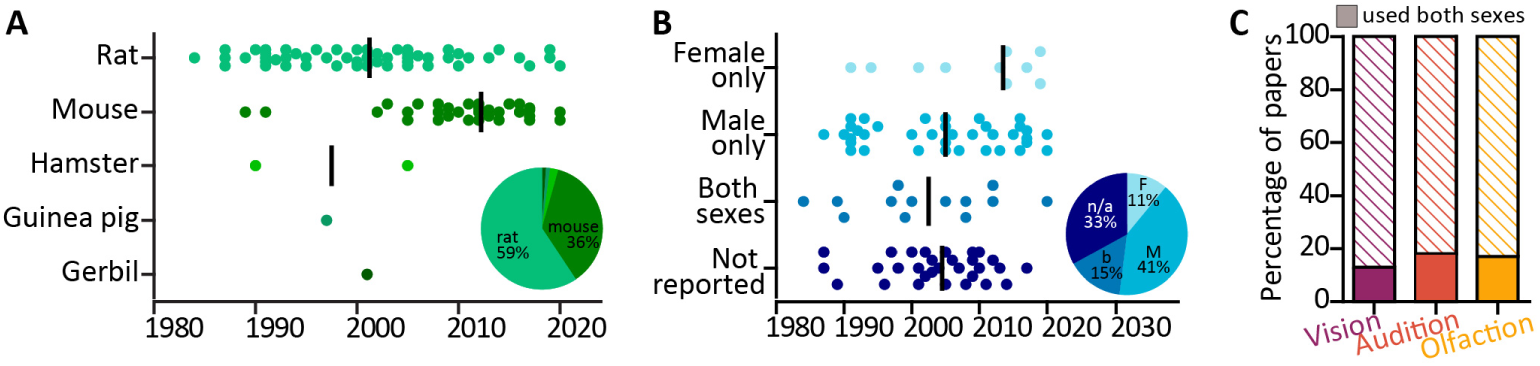
Animal models. (A) Distribution of rodent species used over time (1984-2021). Dots are individual studies, line indicates median publication year. Insert: pie chart reporting the percentages of the 92 selected studies using rats (59%), mice (36%), and other rodents (5% split among hamsters, guinea pigs and gerbils). **(B)** Distribution of rodent sex used over time (1984-2021). Dots are individual studies, line indicates median publication year. Insert: pie chart reporting the percentages of the 92 selected studies using females only (11%), males only (41%), both sexes (15%), or failing to report the sex of the used animals (33%). **(C)** Proportion of papers using both sexes (filled rectangles) across the three sensory modalities: purple, vision = 13%; orange, audition = 18%; yellow, olfaction = 17%. Striped rectangles include studies which used only one sex, or failed to report the sex used.

### Features of the experimental paradigm: diverse methods to induce sensory deprivation

Next, we focused on how visual, auditory, and olfactory deprivations were induced. The most common method to induce sensory deprivation involves lesioning the peripheral sensory organ via surgical or chemical approaches (surgical lesion: n=63; chemical lesion: n=13; lesion methods combined = 82%; Fig. 5A). Other less invasive methods were sense-specific, and included using nose and ear plugs for, respectively, olfactory and auditory deprivation, and dark rearing for visual deprivation (other methods: n=17, 18%; Fig. 5A). The minimum duration of deprivation significantly differed among sensory modalities (one-way ANOVA; F (2, 87) =5.13; p=0.0078), with the audition field adopting significantly shorter deprivation durations (4.9 days ± 5.7 days) compared to olfaction (18.05 days ± 15.65 days), and a trend of shorter duration compared to vision (13.16 days ± 2.791 days,; Tukey’s post-hoc audition vs olfaction p*<*0.01, audition vs vision p=0.09, vision vs olfaction p=0.36; Fig. 5B). We also assessed how many studies used a reversible deprivation method which allows investigating the circuit recovery while leaving it anatomically intact. Such methods include nose and ear plugs, and eye patches or dark rearing. We found that, compared to vision and audition, a larger fraction of olfactory papers used reversible methods, chiefly the insertion and removal of a nose plug (vision=26%, audition=18%, olfaction=90%: chi-squared test; X2(2)=126; p*<*0.0001; Fig. 5C). Consequently, olfactory and auditory studies often investigated the functional recovery after cessation of the sensory deprivation (vision=3%, audition=18%, olfaction=17%: chi-squared test; X2(2)=12.72; p=0.0017; Fig. 5D). It is noteworthy that while not every olfactory study using nose plugs investigated functional recovery, every auditory paper employing reversible deprivation techniques, *i.e.* ear plugs, studied the circuits recovery after plug removal.

**Figure 5.**
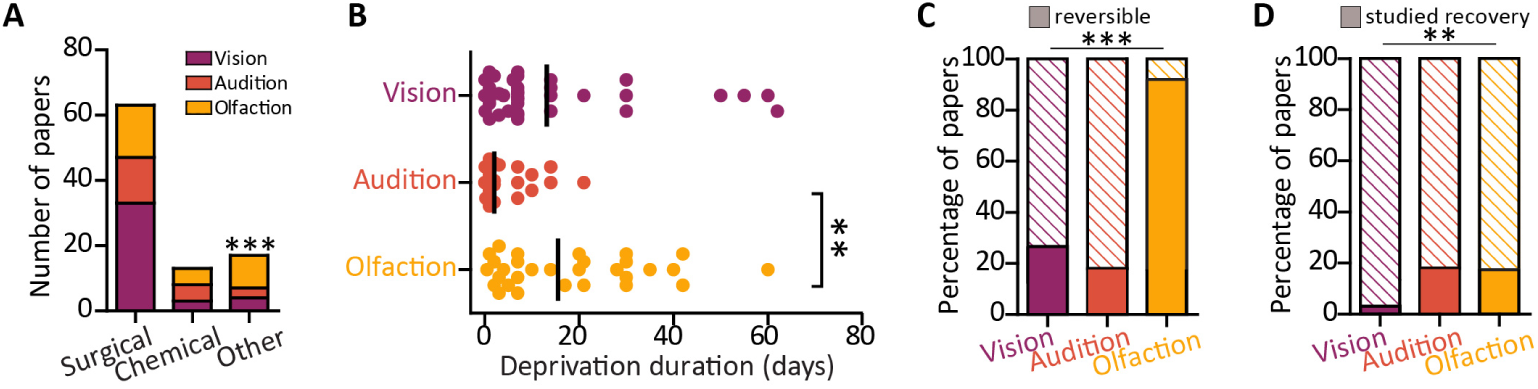
Deprivation method. (A) Number of papers using surgical, chemical, or other (plugs, patches) deprivation methods. **(B)** Minimum deprivation duration in days used in each study (individual dots) and mean duration (black line). **(C)** Proportion of papers using reversible deprivation methods (filled rectangles) across the three sensory modalities: purple, vision = 26%; orange, audition = 18%; yellow, olfaction = 90%. **(D)** Proportion of papers which used a reversible method to induce deprivation and investigated recovery (filled rectangles) across the three sensory modalities: purple, vision = 3%; orange, audition = 18%; yellow, olfaction = 17%. ** p*<*0.01, *** p*<*0.001

### Features of experimental paradigm: deprivation-induced plasticity read-outs

After confirming that sensory deprivation is induced in two broadly similar ways across the three sensory modalities – permanent lesions or reversible removal of the sensory stimuli - we proceeded to assess how the consequences of such sensory deprivations were investigated. Across all sensory modalities, the predominantly employed experimental technique was histology (n=75), followed by molecular biology assays (n=14) and electrophysiological recordings (n=14), and functional imaging in live tissue (n=13). Although histology was the most commonly used technique, the proportion of studies employing each technique is different across the senses (chi-squared test; X2(8)=38.34; p*<*0.0001). Notably, autoradiography is used almost exclusively by vision researchers (except for 1 auditory study; n=7, Fig. 6A). We observed more pronounced differences when we analysed the number of studies employing more than one method to probe deprivation-induced plasticity in the target circuits. While approximately a quarter of papers focussing on visual and auditory early brain areas used more than one technique, over half of olfactory papers investigated the consequences of smell deprivation in the olfactory bulb using multiple methods (vision=26%, audition=23%, olfaction=60%: chi-squared test; X2(2)=36.51; p*<*0.0001; Fig. 6B).

**Figure 6.**
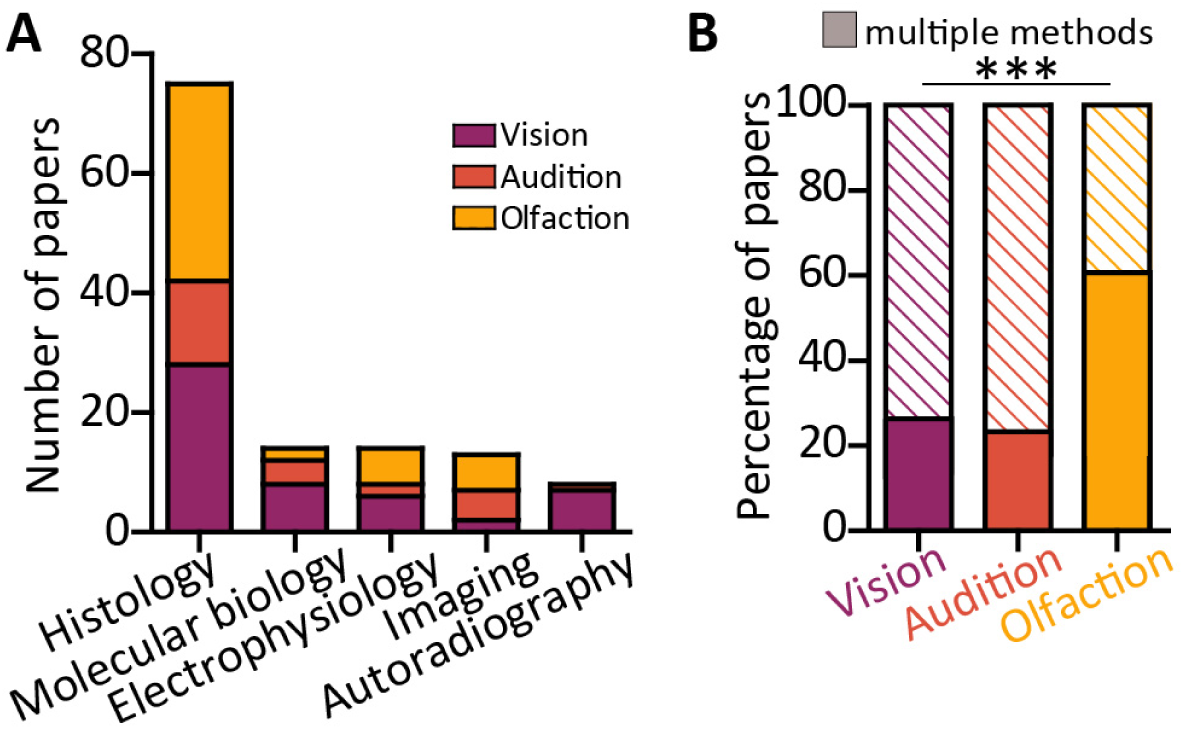
Plasticity readouts. (A) Number of papers across the three sensory modalities using each investigative methodology. **(B)** Proportion of papers using more than one method to investigate the effects of deprivation (filled rectangles) across the three sensory modalities: purple, vision = 26%; orange, audition = 23%; yellow, olfaction = 60%. ***=p*<*0.001

### Features of experimental paradigm: definition of cell types

As experience-dependent plasticity is cell-type specific(23,30), we next assessed on which cell types the 91 selected studies focussed their investigation. We broadly grouped cell types into glia and neurons, and further subdivided the neuronal class into principal neurons, whose axon projects out of the circuits where their soma resides (*i.e*, relay neurons in the lateral geniculate nucleus and superior colliculus, bushy and T-stellate cells in the ventral cochlear nucleus, fusiform and giant cells in dorsal cochlear nucleus, mitral/tufted cells in the olfactory bulb), and excitatory or inhibitory interneurons, whose processes are fully contained in the local circuit. We found that a significant number of studies across all three sensory modalities, but more pronouncedly in vision, did not define the cell types that they investigated, either because they looked at area-wide measures or because they did not classify they types/subtypes of cells that they assessed (number of studies defining cell types: vision=13%, audition=59%, olfaction=80%: chi-squared test; X2(2)=93.96; p*<*0.0001; Fig. 7A). Given the historical popularity of visual deprivation (Fig. 3C; mean publication year 2002), we analysed the trend over time for studies to define cell type (Fig. 6B). As the median year for cell type definition is 2009, we confirmed that defining cell types has become more routine in recent years (*e.g*., the two most recent papers in vision both defined cell types(31,32), but not fully penetrant since long-established practices in each field seem to somewhat linger. When papers defined the cell types in which they investigated deprivation-induced plasticity, they did so in different proportion across the three sensory modalities. While principal neurons have been explicitly investigated in all circuits albeit with different proportions (2/39 papers in vision(31,32), 3/30 papers in olfaction(33–35), and 7/22 papers in audition(36–42)), a much more diverse picture emerges when one focuses on of interneurons. Notably, while none of the vision papers investigated interneurons explicitly, most of the olfactory studies did, both historically and currently (Fig. 7C-D). Studies investigating the olfactory bulb focussed on both inhibitory interneurons (23 papers, 77% of all olfaction papers) and excitatory interneurons (5 papers, 17% of all olfaction papers; see Table 1 for details). Similarly, in audition, a fair percentage of papers investigated inhibitory interneurons (5 papers, 23%), and excitatory interneurons (2 papers, 9%). Such difference was expected given the different ratios of interneurons present in these circuits, with the OB totalling over 80% of neurons being GABAergic(17). In recent years more attention has been devoted to glial cells in all circuits (median year of investigation =2003).

**Figure 7.**
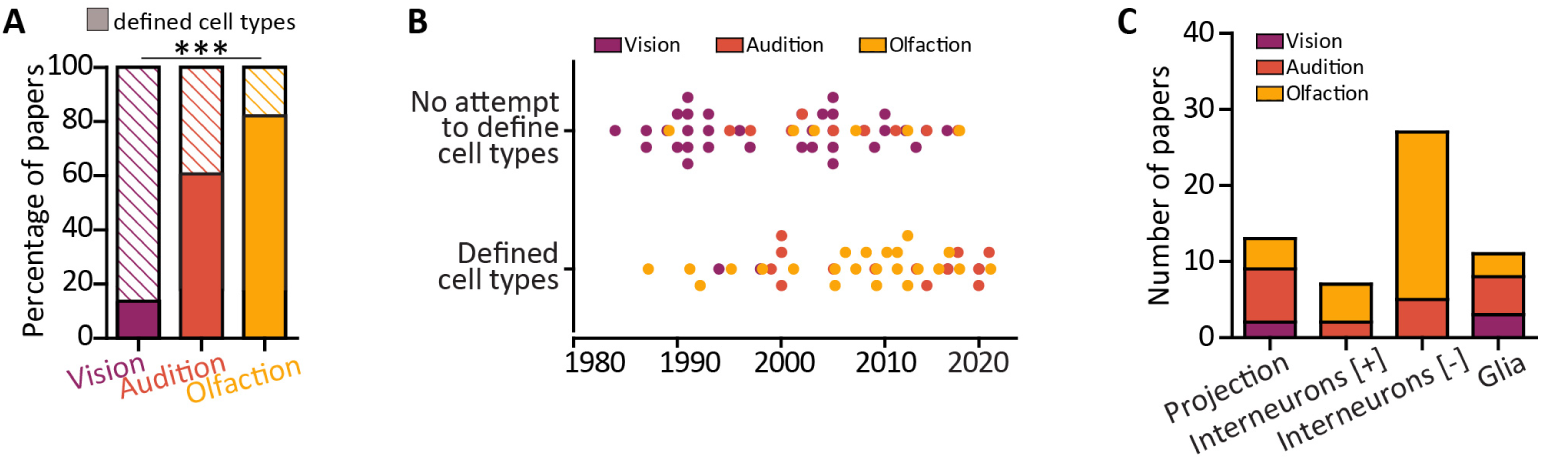
Definition of cell types. (A)Proportion of papers defining the cell types where the deprivation-induced plasticity was investigated (filled rectangles). *** p*<*0.001 **(B)** Distribution of papers over time split by the lack/presence of cell types definition. Dots are individual studies; purple = vision, orange = audition, yellow = olfaction. **(C)** Among the studies which defined cell types, number of papers investigating deprivation-induced plasticity in projection/principal neurons, excitatory [+] or inhibitory [-] interneurons, and glia across the three sensory modalities. Note the lack of vision papers focussing on interneurons.

**Figure 8.**
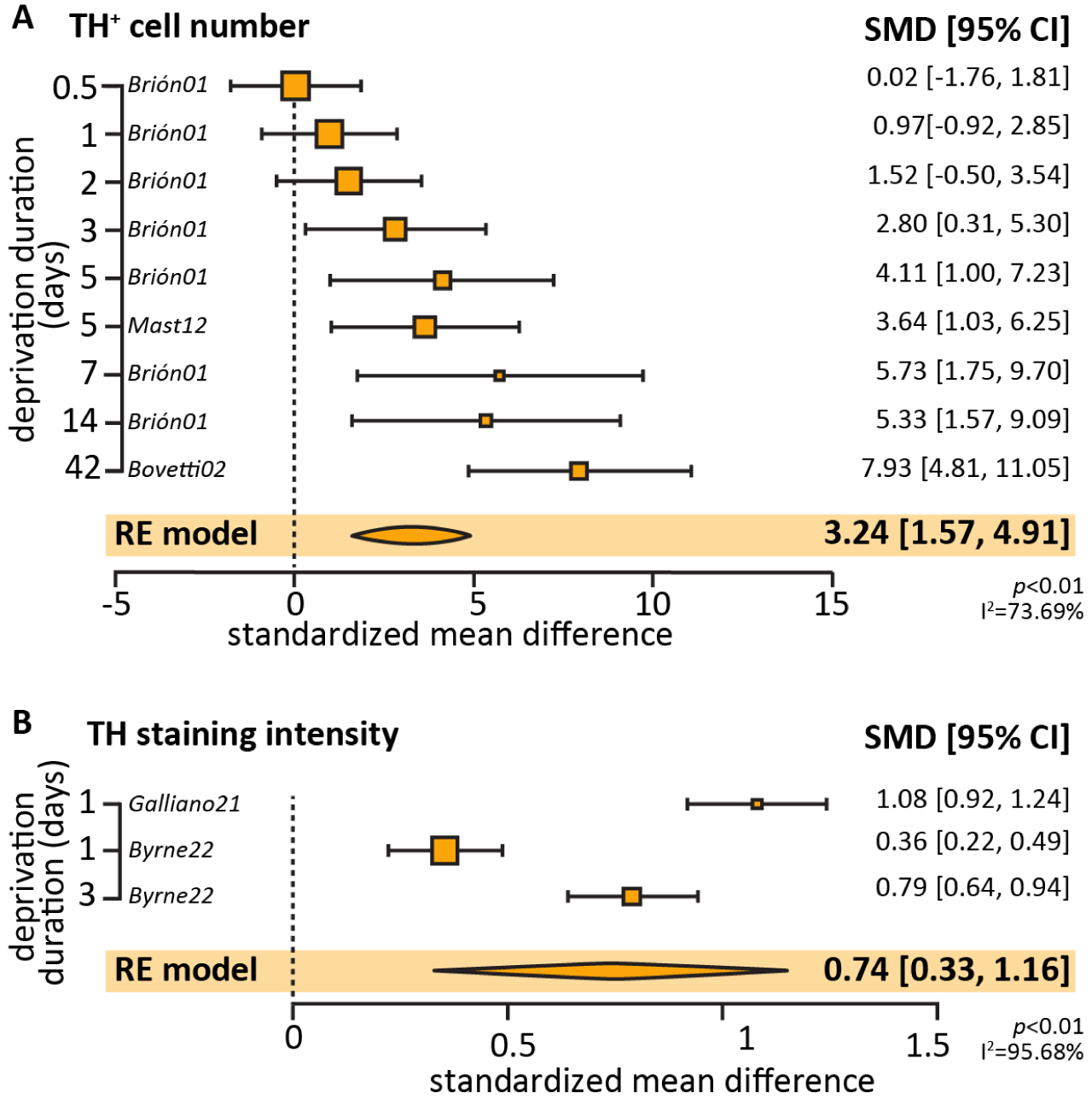
Metanalysis of TH expression after olfactory deprivation of various durations. (A) Effect size of olfactory deprivation on the number of TH-positive DA neurons in the OB. Note that 7/11 datapoints originated from the same study(570). **(B)** Effect size of olfactory deprivation duration on the TH staining intensity in bulbar DA neurons. Note that 2/3 datapoints originated from the same study(120).

### Effects of sensory deprivation-induced plasticity

Our objective was to assess the consistency, directionality, and comparability of the deprivation-induced effects across systems in a quantitative manner, performing a meta-analysis of the literature. Unfortunately, this proved impossible given the huge variability in the deprivation-induction method (Fig. 5), experimental techniques employed to readout the deprivation-induced plasticity (Fig. 6), as well as severity of deprivation (*i.e.* lesion or reversible stimulus removal, Fig. 7) and completeness of the reporting. Reliable activity-dependent molecular markers can be used to validate the success of a deprivation method. However, in vision and audition such a pronounced and stereotypical change has not been systematically described, albeit calcium binding proteins and immediate early genes are potential candidates(43–45). In olfaction, reduced expression of tyrosine hydroxylase (TH) has been traditionally used to confirm that the nose plugging or cauterization had an effect(46). As such, TH RNA and/or protein expression is the most reported plasticity readout across the literature. Using the same search terms to extend the publication date to March 2023 to increase our paper pool to a statistically-meaningful size(47), we collated 11 papers reporting TH changes using immunofluorescence and we attempted to conduct a meta-analysis to assess the overall effect of olfactory deprivation on TH expression. While all studies showed a decrease in TH, we wanted to see if the change was dependent or modulated by deprivation duration. Out of the 11 papers, the ones with sufficient statistical reporting allowing for meta-analysis (5 papers) were split into ones that used TH-positive cell density (9 experiments across 3 papers) vs. those based on TH cell fluorescence (3 experiments across 2 papers) to report TH changes. The overall measurement of standardised mean difference was significant for both measurements, 3.24 (random-effects meta-analysis, p = 0.0001) for density and 0.74 (random-effects meta-analysis, p = 0.0004) for the staining intensity, demonstrating that olfactory deprivation is correlated with a decline in TH expression. However, upon further analysis, the test for heterogeneity in each case (I^2^ = 73.69%, p = 0.0003 and I^2^ = 95.68%, p *<*0.0001) shows that we cannot assume that there is a singular effect size for the population represented by these studies. When occlusion duration was added as a modulator, the amount of heterogeneity unaccounted for by occlusion time was very high in studies investigating TH by fluorescence. This was lower for studies of TH density analysis (I^2^ = 23.07%, p = 0.2486), suggesting a more homogeneous underlying effect size once occlusion duration was accounted for. The low number of experiments and papers entered in the meta-analysis precludes any conclusions from being drawn as the number of studies falls below the threshold of minimum 10 experiments used by meta-analyses in the field(47).

Given the impossibility to perform quantitative meta-analysis on the entire dataset, we attempted to summarise and correlate these results qualitatively by grouping the findings using the only descriptor present in every study: the overall duration of the deprivation, which we divided in short-term (1 day or less), medium-term (up to a week), and long-term (over a week). This grouping, albeit arbitrary, reflects physiologically relevant scenarios of changes in sensory inputs. Short-term deprivation reflected transient and mild diseases involving sensory organs, for example, temporary hearing loss from noise overexposure(48), loss of smell from mild colds(49), and transient visual loss(50). Medium-term deprivation reflects longer albeit still temporary diseases, such as acute ear infection(2) and anosmia following COVID-19(51). Long-term deprivation reflects severe, more permanent forms of sensory deprivation, such as presbycusis (age-related hearing loss(52)), long COVID-19(53) and diabetic retinopathy(54). Where possible we tried to group findings by cell-type (Fig. 7) and broad subregion (*e.g.* dorsal LGN/CN vs ventral LGN/CN). When cell type specificity was not reported, the results were divided into a few broad themes, namely, macroscopic size, protein expression, activity, and proliferation. Using this structure a few central commonalties emerge and are summarized below and in Table 1.

Given the impossibility to perform quantitative meta-analysis on the entire dataset, we attempted to summarise and correlate these results qualitatively by grouping the findings using the only descriptor present in every study: the overall duration of the deprivation, which we divided in short-term (1 day or less), medium-term (up to a week), and long-term (over a week). This grouping, albeit arbitrary, reflects physiologically relevant scenarios of changes in sensory inputs. Short-term deprivation reflected transient and mild diseases involving sensory organs, for example, temporary hearing loss from noise overexposure(41), loss of smell from mild colds(42), and transient visual loss(43). Medium-term deprivation reflects longer albeit still temporary diseases, such as acute ear infection(2) and anosmia following COVID-19(44). Long-term deprivation reflects severe, more permanent forms of sensory deprivation, such as presbycusis (age-related hearing loss(45)), long COVID-19(46) and diabetic retinopathy(47). Where possible we tried to group findings by cell-type (Fig. 7) and broad subregion (*e.g.* dorsal LGN/CN vs ventral LGN/CN). When cell type specificity was not reported, the results were divided into a few broad themes, namely, macroscopic size, protein expression, activity, and proliferation. Using this structure a few central commonalties emerge and are summarized below and in Table 1.

### Common mechanisms of plasticity: short-term deprivation produces decreased activity and disinhibition

After short-term deprivation, defined as 24 hours or less, the effect most consistently investigated across all three senses was changes in overall activity levels. Despite using different proxies, a reduction in neuronal metabolic activity in the early brain circuits receiving input from the deprived eye/ear/nostril was found across all studies. In the visual system, this is manifested as a reduction in glucose uptake in the LGN and SC(55,56). Similarly, a decrease in the immunoreactivity of the activity early gene cFos, was observed in the olfactory(33,57) and auditory systems(58). This rapid decrease in activity is in alignment with that found in primary visual and auditory cortices(18,52).

Deprivation-induced homeostasis can be achieved, at least transiently, by balancing the changes in the excitatory and inhibitory pathways, and it has been proposed that rapid disinhibition mediated by downregulated inhibitory networks precedes excitatory plasticity(59,60). The papers that defined cell types and specifically investigated inhibition reported findings which were consistent with decreased inhibition: GABAergic and glycinergic mechanisms are downregulated following brief sensory deprivation (see Table 1 for details). In the auditory system, GlyR1 postsynaptic density in bushy and fusiform cells was decreased(42). In the olfactory system, the dopaminergic inhibitory interneurons downregulate TH expression, reduce their intrinsic excitability, and shorten the axon initial segment(33). In visual areas, immunoreactivity of GABA transporters (GAT-1 and GAT-3) in hypertrophied astrocytes in the de-afferented SC was increased, suggesting an increased uptake of GABA(61). This early disinhibition of the deprived system, achieved via different mechanism in each circuit, also occurs in higher areas and has been extensively reviewed in the auditory, visual and somatosensory cortices(52,60,62).

Conversely, alterations in excitatory signalling following one day-long deprivation were less clear cut. In olfaction and vision, studies which investigated glutamatergic function found no differences after brief deprivation. In vision, this was investigated using autoradiography, and no changes were found in the SC nor dLGN for AMPA, NMDA and kainate receptors expression(63). In the olfactory system, while both the glutamatergic principal neurons and interneurons downregulated the expression of immediate early genes and activity markers, they did not modulate their intrinsic excitability nor their axon initial segment morphology(33). In the auditory system, however, there is a highly specific upregulation of AMPA receptor subunits at the auditory nerve to principal neuron synapse (GluR3 upregulated at bushy cell and fusiform synapse, GluR4 downregulated at fusiform synapse)(42). When combined with concomitant changes in inhibition, the overall functional effect of these changes on circuit homeostasis remains to be elucidated. In contrast to the other two senses, papers in audition mostly focused on short deprivation, with almost half of all papers investigating 1 day or less than 1 day deprivations (9/22; Figure 5B) and glial responses. An increase in glial activation was found across different studies using varied experimental methods and included increases in immune related genes(64), activation(64,65) and proliferation(66) of microglia, as well as increases in staining for astrocyte markers(65). These findings are largely consistent with an increased glial activity in the cochlear nuclei but given the lack of similar investigations in vision and olfaction, it remains unclear whether this is a unique feature of the early auditory circuits or, like decrease in activity and disinhibition, a common response to brief sensory deprivation.

### Common mechanisms of plasticity: medium-term deprivation is reflected by an increased involvement of glia and synaptic remodelling

After mid-term deprivation, defined as lasting between one day and a week, alongside continued decreases in activity the most prominent changes were changes in synaptic properties and glial activation. Across all three sensory modalities, multiple studies reported functional and structural changes at the synapses formed by the cranial nerve axon terminals, suggesting an overall different circuit-level excitation/inhibition (E/I) balance. In the olfactory system studies, investigations did not focus on presynaptic remodelling but rather on the postsynaptic properties of the various interneuron subtypes innervated by the olfactory nerve directly or via multi-synapses loops. Functional changes have been found at the excitatory interneurons (external tufted cells, ETCs), where deprivation decreased the amplitude of spontaneous inhibitory postsynaptic currents(67), as well as increased quantal glutamatergic (AMPA-mediated) postsynaptic currents(68). Structurally, a reduction of spine density was found in the inhibitory interneurons granule cells(69) as well as GAD67 puncta in ETCs(67). Since ETCs modulate both inhibitory interneurons as well as excitatory principal neurons in the OB, the overall effect of these changes on the whole-circuit output remains unclear but seems to suggest modifications in the E/I balance.

In the visual system only synapses formed by the optic nerve synapses onto principal neurons in the dorsal LGN were studied after medium-term sensory deprivation. Thalamocortical neurons in the dLGN had smaller single-fibre AMPA-mediated postsynaptic currents at the retinogeniculate synapse(70,71), indicating weakening of individual glutamatergic afferents following deprivation. However, this was counterbalanced by a concomitant increase in excitatory postsynaptic current (EPSC) amplitudes, indicating an increase in the number and/or strength of retinogeniculate synapses(70). Structurally, there was an increase in the phosphorylation of stargazin, a transmembrane AMPAR trafficking protein in the LGN involved in synaptic scaling(72). Overall, the evidence indicates that visual deprivation induces synaptic remodelling at the excitatory retino-geniculate synapse in the dLGN after medium term deprivation. In the auditory system, the most prominent synaptic effect of one week deprivation is the overall macroscopic decrease in the number of synaptic contact zones in the anterior VCN(73). Along similar lines, a significant reduction in VGLUT1 staining was found in the VCN(74). Interestingly, inhibitory synapses decreased in number less markedly than excitatory ones(73). Taken together with the findings of rapid disinhibition described above, this could suggest a period of transient over-inhibition following an early disinhibition phase of the VCN after cochlear deafferentation.

In addition to changes in synaptic transmission, an increase in glial proliferation and activation was found consistently across all senses. In the olfactory system, significant increases in the density and activation of microglia are seen in all papers which investigated this phenomenon(69,75). The increased activation of microglia is evident both in the morphology (shown as fewer primary microglial processes and shift towards hypertrophied morphology)(75) and immunoreactivity staining with specific marker for activated microglia CD68(69). Moreover, a significant increase in reactive astrocytes was reported in all OB layers(76), as well as in the visual system (although here it was not a formal strand of investigation)(77). In the auditory system in addition to increases in microglial and astrocytic immunoreactivity(65,78), glial activation has been linked to synaptogenesis in both the VCN and DCN(74,79). During the first three days of deprivation, astrocytes increase production of neurocan, aggrecan, and MMP9, which are key for synapse stabilisation (neurocan and aggrecan)(80,81) and the induction of synaptic plasticity via ECM remodelling (MMP9)(82). Then, between three and seven days of deprivation, expression of these proteins decreases and is replaced by PSA-NCAM and MMP2. Both are strongly associated with structural plasticity, MMP2 via neurocan digestion and PSA-NCAM by mediating synaptogenic interactions between neurons and astrocytes(83,84). This astrocytic biphasic response and its potential role in synaptic remodelling warrants further research in the visual and olfactory regions.

### Common mechanisms of plasticity: long-term deprivation

Given the adopted definition of long-term deprivation as any period spanning between a week and a year, we found that the effect on experience-dependent plasticity on the different cell types was more heterogeneous than for shorter-lasting deprivations. Variability notwithstanding, remarkably consistent macroscopic changes and apoptosis were found across all three senses. In olfaction, this is reflected by smaller overall OB volume and weight(85–87) as well as in a reduction of external plexiform layer thickness(85) and glomeruli volume(88), and in increased cell death in the mitral and granule cell layers(89). Similarly, increased apoptosis was seen in the dLGN(90) and reduction in volume was seen in the stratum zonale, stratum griseum superficiale and straum opticum regions of the SC(91,92). In audition, there was a marked decrease in VCN area and neuron quantity(93). This contrasts with short-term deprivation, where macroscopic structural changes were scarcely investigated. To this date, only one paper reported no difference in apoptotic neurons between the unperturbed and deprived side(94). In the olfactory system these macroscopic changes can manifest in a rather unique way, namely, in cellular turnover. Neural progenitors are produced in the subventricular zone and migrate along the rostral migratory stream (RMS). They arrive at the subependymal layer of the OB, where they differentiate into glomerular interneurons or granule cells(95). The experience-dependent nature of adult neurogenesis has been widely researched (reviewed in (96)), and it is well known how olfactory deprivation and enrichment can decrease or increase the survival of adult born neurons, respectively. The findings of this review were in alignment with the literature, where after medium to long term deprivation, neurogenesis is reduced at each level of the neurogenic process. After 2-3 weeks deprivation, there was a reduction in the integration of neuroblasts into the OB, as well as slower migration along the RMS(87). In the glomerular layer, a decrease of newborn periglomerular cells was found after 28 days(97) and 42 days(98). This is mirrored by an increase in cell apoptosis in both the periglomerular(97) and granule cells(99). Remarkably, this experience-dependent neurogenesis is found to be very specific to the dopaminergic population at the glomerular layer, as the calbindin and calretinin glomerular interneuron populations remain unchanged after deprivation(97). Unfortunately, no papers investigated neurogenesis in the context of shorter deprivations, so we cannot correlate these findings with the deprivation duration.

According to theoretical frameworks and working models in the homeostatic plasticity field, the expectation is to see prominent changes in excitatory networks with a long-lasting sensory deprivation as the circuit stabilizes to a new set-point(60). While no papers focused on the early visual circuits’ principal neurons responses to long term deprivation, changes at the excitatory synapse between incoming nerve and principal neurons were seen at bushy cells in the cochlear nuclei. Ten days of auditory deprivation using earplugs was correlated with a decrease of the presynaptic marker VGlut immunoreactivity and size of synaptic vesicles(38). A similar VGlut decrease was observed in the VCN 3-14 days after cochlear ablation, suggesting a robust effect in both manipulations(74). Postsynaptically, GluA2/3 expression was upregulated, while GluA2 and GluA4 remained unchanged(38) - an effect found also after 1 day deprivation(42). In the olfactory system, the systematic search only returned one paper which investigated mitral/tufted cells after long term deprivation(34). After 60 days deprivation, while there were no changes in their spontaneous activity rates, the number of mitral/tufted cells responding to more than one odour was increased. In addition, significantly more cells in the deprived bulb responded to specific odours presented at higher intensities, suggesting a possible decrease in odour discrimination coupled with increase in responsiveness. Conversely, in a very recent paper published after the cut-off date of our systematic search(100), the authors found a shortening in the mitral cell axon initial segment as well as spiking frequency after unilateral naris occlusion for 30 days compared to the un-occluded bulb, indicating potential for decrease in intrinsic excitability at OB principal neurons after longer deprivation times. The slightly discrepancy in mitral/tufted cell firing rate after long deprivation found by(100)and(34) could be attributable to various methodological differences, such as occlusion times (30 days vs. 60 days, respectively), as well as experimental methodology (*in vivo* electrophysiology vs. acute slice electrophysiology, respectively). In addition to changes at the principal neurons, deprivation on a longer time scale also sees a decrease in glial activation, which was investigated in vision and audition. In vision, the number of immunoreactive astrocytes significantly decrease 12-48 weeks after enucleation(101). In audition, the number of calbindin positive astrocytes starts to decrease 30 days after cochlear lesion(78).

## Discussion

In this study we employed systematic and meta-analysis methods to investigate the effects of sensory deprivation of various durations at the homologous regions receiving the cranial nerve input of three sensory modalities - vision, olfaction and audition. Our analysis returned large disparities across sensory modalities in publication trends, experimental methodologies employed, as well as focus of the research. However, despite such methodological and reporting differences, a few shared findings describing adaptive responses and compensatory mechanisms to deprivation can be extrapolated.

### Profound differences in micro circuitry, experimental protocols, plasticity readouts, and reporting make comparisons across the three sensory modalities challenging

The three senses differ in their micro circuitry, as well as in the degree to which it has been characterised (Fig 1). The lack of studies defining cell types and investigating interneurons in vision could be partially attributed to the incomplete characterisation of the LGN, where cell identification challenging by its lack of overt lamination(20). Indeed, while the OB is a highly inhibitory circuit where the ratio of GABAergic interneurons to excitatory neurons is much higher than other parts of the brain(24), visual structures such as the LGN are comprised of mainly excitatory projection neurons(102).

The main experimental methods used to induce and interrogate plasticity also differed widely across the three senses. While all sensory deprivation approaches are long-established(103,104), visual deprivation via monocular enucleation and eyelid suturing garnered prominence and widespread adoption after the seminal work of Hubel and Wiesel(17). Thus, many vision papers in our list were older and tended to use this surgical technique, while papers focussing on olfactory and audition deprivation are more recent and more commonly employ the well-established reversible procedure of nose and ear plugging.

Of note, contrary to rod, cones, and hair cells, the olfactory sensory neurons are capable of regenerating throughout the life of the animal(105). This poses an important difference among senses. While deprivation by deafferentation (direct lesion to the peripheral sensory neurons) and by sensory deprivation (removal of sensory stimulation while leaving the anatomy of the circuit intact) has largely similar effects in the visual and auditory systems as both are incapable of regenerating, the effects in the olfactory system can be divided into adaptive plasticity (elicited by sensory deprivation, *e.g.* nose plug, and within the remit of this study) and regenerative plasticity (elicited by ablation of OSNs using olfactotoxic drugs such as methimazole and dichlobenil)(106,107). Adding onto the anatomical and procedural discrepancies across the senses, a profound lack of consistency in reporting precluded the possibility of a systematic meta-analysis. Furthermore, the findings of this study are heavily subject to publication bias, as the lack of findings in a particular area does not necessarily denote a negative finding. For example, we found an abundance of studies demonstrating the role of glia in regulating the cochlear nuclei response to short auditory deprivation, but it remains unclear whether this is because glial plasticity was never investigated olfaction and vision, or it was investigated, found absent, and not published.

### Despite procedural differences, the three early sensory areas share deprivation-dependent plasticity motifs

With the caveat that the short-medium-long deprivation duration classification is partially arbitrary, albeit based on clinical evidence(2,48–54), we found four consistent plasticity responses across the senses (Fig. 9). First, for sensory deprivation lasting up to 24 hours, the overall neuronal metabolic activity is consistently reduced. In addition, the dampening of inhibitory interneuron activity was consistently reported in olfaction and audition (but not investigated in vision), suggesting fast-acting circuit disinhibition. This is in line with data from sensory cortices and the working hypothesis that excitatory principal neuron plasticity is preceded by a depression of inhibitory interneurons(59,60). Second, in these early sensory areas, long-term and/or permanent sensory deprivations result in more drastic changes involving macroscopic changes to the circuit architecture which manifests as tissue shrinkage, cell apoptosis, and reduced adult neurogenesis in the olfactory system. Third, there is consistent evidence across all three early sensory areas and all deprivation durations for glial activation and proliferation. This highlights the prominent role of glia in shaping neuronal plasticity(13). Fourth, although less clear-cut, from medium-term sensory loss onwards there is a strong indication of synaptic remodelling and changes in E/I balance. The fact that these four effects were revealed from the systematic search despite profound differences in circuit architecture, methodology, and readouts is perhaps an indication that these plasticity motifs are robust and conserved.

**Figure 9.**
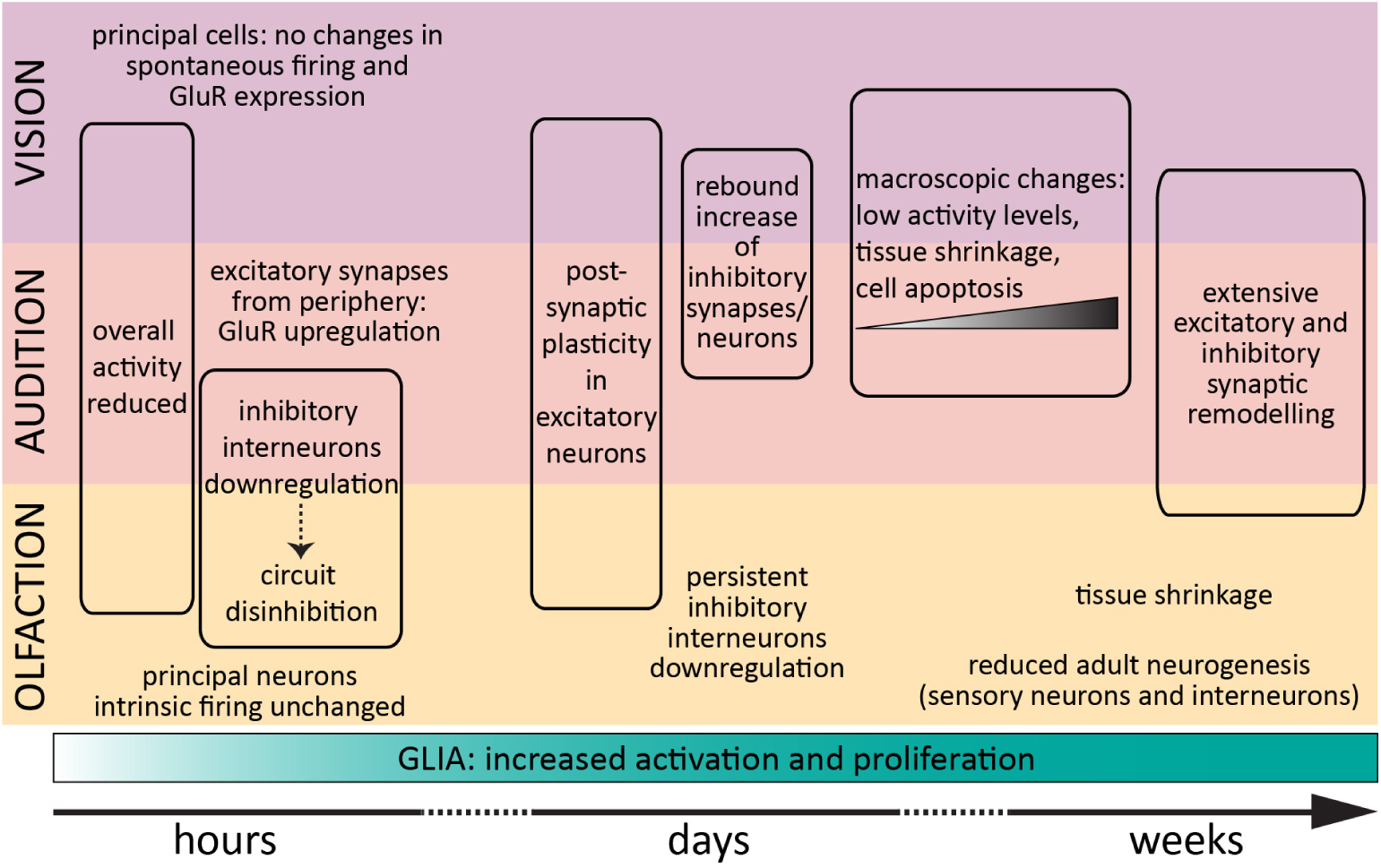
Deprivation-induced plasticity. Graphical representation of the cellular and circuit effects of sensory deprivations of increasing durations in early visual, auditory, and olfactory areas. See Table 1 for details and references.

It remains unclear whether these findings can be generalized to other areas. The time-course and multiplicity of mechanisms of deprivation-induced plasticity in pre-cortical regions found in this study align with those found in cortex and fit with the canonical theories of homeostatic plasticity(10,11). Notably, the phenomenon that the inhibitory cells are modulated rapidly after sensory deprivation and that such changes precede adaptations in excitatory neurons has been widely reported in sensory cortices. For example, after 1 day of monocular deprivation parvalbumin-positive GABAergic basket cells in the primary visual cortex significantly reduced their firing rate and receive less synaptic excitation, whilst pyramidal cells remained unchanged(108,109). This rapid functional reduction of inhibitory tone is also supported by structural changes(110), and similar dynamics have been described in primary auditory(111–113) and somatosensory (62,114) cortices. Interestingly, the opposite manipulation – 6h sensory enrichment via whisker stimulation - has been shown to induce rapid downregulation of barrel cortex pyramidal cells’ excitability(115), perhaps a neuroprotective mechanism to combat runaway excitation. This highlights a possible asymmetry in plasticity induced by sensory deprivation vs. enrichment, which warrants further investigation in both cortical and subcortical regions(10).

Finally, it has been suggested that rapid disinhibition restores the system to a “juvenile” state of plasticity(15,16) and facilitates functional recovery by creating an environment more permissive for the induction of synaptic potentiation through long-term potentiation or spike-timing dependent plasticity in excitatory neurons(16,62,116,117). However, it remains unclear whether the subcortical disinhibition we described reflects the start of functional recovery or simply a response to reduced input(110). This is of particular relevance for those wishing to use data from animal model deprivation studies to inform translational interventions in patients suffering from sudden sensory loss. Further research is required to better dissociate the mechanisms of the two responses, and to understand the relationship between subcortical plasticity and its feedforward consequences in cortex.

### Recommendations

This work synthesized studies on the early stages of sensory processing in audition, vision and olfaction, and the results revealed significant heterogeneity in deprivation method, experimental readouts and reporting, including definition of investigated cell types. While biological restrictions of the circuit (*e.g.* accessibility, different animal models) may explain some of the heterogeneity, this review also highlights areas needing further research. To improve comparability within and between fields, the identification of a ‘gold-standard’ deprivation methods and validation readouts, together with systematic reporting of statistics and deposition of raw data, are sorely needed. Furthermore, the field could improve on better defining cell-types, especially with increasing evidence, including from our results, that inhibition and excitation are differentially regulated after deprivation(11,118,119). Finally, since most published work focused on deprivation-induced plasticity in primary sensory cortices, it is essential to fully characterize potential effects in the pre-cortical areas which link them to the periphery. As described above, these are not passive relays but active plasticity hotspots which send to cortex already processed and modulated inputs which must be considered to depict the full picture of the final cortical computation and behavioural outputs.

## Conclusions

In conclusion, this systematic review and literature meta-analysis found that, notwithstanding major experimental heterogeneity in inducing and assessing deprivation-induced plasticity, the early visual, auditory, and olfactory systems largely employ shared mechanisms to change their circuit processing. Future work should strive to standardise both experimental and reporting approaches and investigate how early circuits and higher cortical areas together coordinate an appropriate adaptive response to a lack of peripheral sensory inputs.

## Methods

### Inclusion criteria

We included papers which met all of the following criteria: (a) rodent animal models (mouse, rat, guinea pig, hamsters, gerbils), (b) primary research, (c) English language, (d) sensory deprivation performed (d’) *in vivo* (d”) in animals adults for the entire duration of the deprivation (postnatal day 21 and over), (e) plasticity investigated at the location of first central synapse after cranial nerve (*i.e*. olfactory bulb, lateral geniculate nucleus, superior colliculus, dorsal and ventral cochlear nuclei, trigeminal nucleus).

### Search Strategy

Two separate searches, without any time constraints on publication date, were performed on November 22nd 2021 in PubMed and Scopus using the Boolean search strings detailed below:

*“ ((mouse) OR (rat) OR (gerbil) OR (guinea pig) OR (rodent)) NOT (cross-modal)” AND ((sensory deprivation) OR (auditory deprivation) OR (auditory deafferentation) OR (visual deprivation) OR (dark exposure) OR (enucleation) OR (retinal lesions) OR (olfactory deprivation) OR (odor deprivation) OR (naris occlusion) OR (nostril occlusion) OR ((trimming) AND (whiskers)) OR ((plucking) AND (whiskers)) OR ((pruning) AND whiskers)) OR (ear plug) OR ((cauterised) AND (naris)) OR ((cauterised) AND (nose)) OR ((cauterised) AND (olfactory)) OR (nose plug) OR ((naris) AND (closure)) OR ((whisker) AND (deprivation))) AND ((superior colliculus) OR (lateral geniculate nucleus) OR (visual thalamus) OR (cochlear nucleus) OR ((trigeminal nucleus) AND (whisker)) OR (olfactory bulb)) AND ((plasticity) OR (adaptation) OR (adaptive) OR (experience-dependent) OR (homeostatic) OR (synaptic scaling) OR (compensatory) OR (activity-dependent plasticity) OR (firing-rate homeostasis) OR (intrinsic excitability))*”

For Scopus, the string was modified by adding “TITLE-ABS-KEY(” at the start, and brace instead of brackets ().

### Study selection

In the first screening stage, the title and abstract of the research papers from both databases were scrutinized by two independent reviewers (LH and FH) to ensure compliance with the inclusion criteria a-c. Duplicates (research papers found in both databases) were removed. Disagreements were resolved by a third reviewer (EG). In the second screening stage, the full text of shortlisted studies was considered against the inclusion criteria d and e.

### Data Extraction

Prior to the search, a list of relevant information to be extracted from each selected paper was agreed upon. These included: (i) publication year, (ii) rodent species, (iii) sex, (iv) age at deprivation onset, (v) deprivation type and its possible reversibility, (vi) method of deprivation, (vii) duration of deprivation, (viii) experimental method used to probe plasticity, (ix) cell type in which the plasticity was investigated, (x) main findings. Papers meeting inclusion criteria were then carefully screened for this information, which was collated in a master spreadsheet.

### Statistical Analysis

Statistical analysis was performed in Prism (Graphpad) and R. Data were checked for normality, one-way ANOVA with Tukey’s post-hoc correction for multiple comparisons was used to assess differences in deprivation duration, Chi-squared tests were used to compare proportions. Significance was set to p*<*0.05. For the meta-analysis, mean and SD as well as n number were extracted manually from the relevant papers, and submitted to meta-analysis in R using the ‘metafor’ and ‘meta’ packages.

## Acknowledgments

We thank Emma Cahill (Bristol) for the inspiration to tackle a meta-research study; Matthew Grubb, Sarah Byford, Ana Dorrego-Rivas (KCL), Marcela Lipovsek (UCL) and Sue Jones (Cambridge) for comments on the manuscripts; and the participants of the KITP “Statistical Learning in the Brain” Programme and the members of the Galliano laboratory for helpful discussions.

---

**Attached Supplementary Table.** Main findings of the included studies.

**TABLE 1.**

| DEPRIVATION DURATION | SENSORY MODALITY | KEY FINDINGS |
| --- | --- | --- |
| Short-term | Olfaction | **Principal neurons:** decrease in immediate early gene pS6 immunoreactivity, no changes in axon initial segment structure nor in intrinsic excitability (33)<br>**Inhibitory interneurons:** decrease in the number of TH+ neurons in dopaminergic cells(57), TH and c-fos immunoreactivity, axon initial segment length, intrinsic excitability(33)<br>**Excitatory interneurons:** decrease in immediate early gene pS6 immunoreactivity, no changes in axon initial segment structure and intrinsic excitability(33) |
|  | Audition | **Principal neurons:** at auditory nerve synapses increased synaptic expression of GluR2/3 (but not GluR2 or GluR4) in bushy cells, increased expression of GluR2/3 and reduced expression of GluR4 in pyramidal cells; no changes at glutamatergic synapses from parallel fibres to pyramidal cells; at inhibitory synapses decrease in the expression of GlyRa1 in bushy and pyramidal cells(42)<br>**Interneurons:** decrease in calretinin immunoreactivity, transient increase in parvalbumin immunoreactivity after 6 and 12 hours but then decrease to control levels by 24 hours (VCN;(37)); no change in GAD65 and vGluT1 staining compared to contralateral side (VCN;(74))<br>**Glia**<br><u>Various:</u> increased immunoreactivity in GFAP and polysialic acid; no change in GAP-43 (or in in matrix metalloproteases MMP-2 (associated re-innervation) and MMP-9 (associated with neurodegeneration) (VCN;(79))<br><u>Astrocytes:</u> increase in astrocyte staining intensity revealed by GFAP (VCN;(65)); increased immunoreactivity in cell membrane- actin cytoskeleton linker ezrin) (VCN;(79))<br><u>Microglia:</u> increase in immune-related genes Ccl12, Csf1 and Cd44 and in activated-microglia marker Iba1 positive cell number (CN;(64)); no change in CD11b staining intensity of microglia but more compact morphology and increase in immunoreactivity for different phosphorylated forms of MAPKs (VCN;(65));increased proliferation of activated microglia (VCN;(66)) |
|  | Vision | **Macroscopic and overall circuit:** no increase in apoptosis (LGN;(94)) reduction in glucose uptake (LGN & SC;(55,56)); no changes in spontaneous activity or spiking temporal patterns (LGN;(121)); no changes in the distribution of glutamate receptors measured with autoradiography (LGN and SC;(63))<br>**Glia:** increased GFAP immunoreactivity (LGN;(122)); increased GAT-1 and GAT-3 immunoreactivity in hypertrophied astrocytes, no change in GABA nor GAD immunoreactivity (SC;(61)) |
| Medium-term | Olfaction | **Macroscopic:** after surgical or chemical ablation, increase in proliferating Ki67-positive OSNs(123) and in proliferating cells in the OB subependymal layer(76); no change in OSN proliferation after nose plugging(123); reduced |
|  |  | olfactory epithelium thickness(124) and smaller OB volume(87); increased immunoreactivity for apoptosis markers caspase-3(125) and TUNEL(76,124) but no evidence for principal cells apoptosis(124)<br>1. **Inhibitory interneurons:** in dopaminergic cells decrease in TH immunoreactivity (57,67,125) and reduction in DA presynaptic terminal density(75); in immature adult-born granule cells reduced spine density(69)<br>2. **Excitatory interneurons:** reduced amplitude of sIPSC and reduced levels of synaptic GAD67 puncta(67), increased amplitude of AMPAR-mediated mEPSCs(68)<br>3. **Glia:** more reactive astrocytosis in all OB layers(76); increase in microglial density(69,75); microglia more dynamic and shifted towards amoeboid morphology(75); greater percentage of neurons wrapped by microglia(75) and increased phagocytosis of adult-born granule cells(69) |
|  | **Audition** | 1. **Macroscopic:** no change in VCN neuron number (VCN;(126)); no change in choline acetyltransferase activity in (VCN & DCN;(127)); lower glucose uptake (VCN & DCN;(128))<br>2. **Excitatory neurons:** reduction of VGLUT1 immunoreactivity (VCN;(74)); reduction in the density of synaptic contact zone stained with GAP-43 (VCN;(129))<br>3. **Inhibitory interneurons:** increase in GAD65 staining (VCN;(74)); increase in the overall fraction of inhibitory synapses and increase of GAP-43-stained nascent synapses (VCN;(73,129)); recovery to control level in calretinin and increase in parvalbumin and calbindin immunoreactivity VCN (37,78)<br>4. **Glia:** increase in GFAP, GAP-43, PSA, and MMP-2 staining intensity (VCN;(65,79)); increase in p-38 and microglial immunoreactivity (CD11b) (VCN;(65)); increase in calbindin-positive astrocytes (VCN;(78)), and increased expression in astrocytes of Ncam and Agg (VCN & DCN;(74)); decrease/plateau of ezrin and MMP-9 (VCN;(79)) and of p-ERK1/2 immunoreactivity (VCN;(65)) |
|  | **Vision** | 1. **Macroscopic:** reduced SC thickness (61); increased number of LGN neurons with darkly stained perikaryon and pyknosis (apoptosis indicator)(77), but no changes detected with ApopTag(94); reduced glucose uptake in LGN(130); in LGN decrease in immediate early genes expression NGFI-A(131) and c-fos(77). No change in cAMP response element binding protein in SC(132), neurokinin-1, GABAa and serotonin-2 binding sites (SC;(133))<br>2. **Excitatory pathways:** increase in maximum AMPAR-mediated EPSC amplitudes and decrease in single-fibre AMPAR-mediated EPSC amplitudes at retinogeniculate synapse (LGN;(70)); increased stargazing phosphorylation at retinogeniculate synapse (LGN;(72)); LGN TS neurons shift responsiveness from monocular to binocular (31)<br>3. **Inhibitory neurons:** weaker calbindin immunoreactivity and increased number of parvalbumin-positive neurons in LGN (77); increased number of calretinin-positive in SC (134)<br>4. **Glia:** increased number of GFAP immunoreactive astrocytes in LGN (77) |
| Long-term | Olfaction | **Macroscopic**<br><u>Proliferation</u>: reduction in Ki67-positive proliferating cells (123), slower migration of neuroblast along the RMS and integration in the OB (87)<br><u>Protein expression</u>: no significant change in expression of neurogranin (postsynaptic calmodulin binding protein (135)<br><u>Size</u>: with nose plug smaller OB volume after 3 weeks (87); no changes in OSN histology after 4 weeks (123); with chemical or surgical ablation decrease in OB weight after 1-6 months (86); smaller OB but no change in cell number after 20 days (85); after 28 days epithelium thickness, OSN histology, apoptosis marker are back to control level (123,124)<br>**Principal neurons**: no change in M/TC number (35); no change in granule cells mediated IPSCs, decreased odour discrimination and lower response threshold, no changes in spontaneous activity (34,136)<br>**Excitatory interneurons**: broadening of ETCs tuning curves (137)<br>**Inhibitory interneurons**<br><u>Various</u>: no change in GAD immunoreactivity after 5-9 weeks deprivation (138), decrease of GAD67 but not GAD65 protein levels 14 days deprivation (139), no change in apoptosis in calbindin and calretinin cells (97); decrease in new-born cell density in the glomerular layer (97,98); in newly generated cells no difference in the expression of GAD67, calretinin, calbindin (98)<br><u>EPL interneurons</u>: no difference in parvalbumin cell number and apoptosis (85,97); reduced GluR1 and GAD65 immunoreactivity (85)<br><u>Dopaminergic cells</u>: decreased TH immunoreactivity and/or decreased TH-positive cell number and/or increased apoptosis (34,45,97,98,138–140); decrease in TH-positive new born cells (98); decrease in dopamine and DOPAC levels, no change to NE or DHPG (45)<br><u>Granule cells</u>: reduction of synaptic puncta in the internal plexiform layer, but not external plexiform layer (88); increased secretagogin expression (140); decreased number and soma size (35); decrease in adult born cell number and significant increase in apoptosis after 4 weeks (99); in newly generated cells reduced density of glutamatergic inputs and of output synapses, while in resident cells increase of synaptic inputs on the proximal dendrites (141) |
|  | Audition | **Macroscopic**<br><u>Size</u>: reduction in VCN area and neuron density (93); decrease in number of GAP-43 nascent synapses (VCN;(129))<br><u>Protein expression</u>: increased choline acetyltransferase activity after 1 month and returns to control levels after 2 months (VCN;(36,127)); increased expression of Kv1.1 and Kv3.1b in the VCN (41); reduction in the fraction of inhibitory synapses but no changes in overall synaptic contact zone in VCN after 70 days (73)<br>**Excitatory pathways**: Decrease in VGLUT1 expression in auditory nerve terminals across both nuclei (93,142); VGLUT2 increases in the interstitial region and fusiform layer of DCN and AVCN (142)<br>**Principal neurons**: decreased glycine immunoreactive puncta in pyramidal cells and bushy cells (36); increased of postsynaptic density thickness and upregulation of GluA3 subunit expression on bushy cells (38) |
|  |  | 4. **Inhibitory neurons:** decreased glycine immunoreactive puncta in both types of VCN stellate cells (36)<br>5. **Glia:** more calbindin-positive astrocytes then after 30 days number starts to decline (VCN;(78)) |
|  | Vision | 1. **Macroscopic**<br>• <u>Size:</u> increase in apoptotic cells number in the dLGN after 12 days but no change in the vLGN (94); decrease in volume and thickness of the superficial layers of the SC (91,92); increased staining intensity of V1 terminals in the upper half of the superficial SC (91); increased density of serotonin immunoreactive fibres in superficial SC and LGN (143); increased density of locus coeruleus to LGN projections if deprived at p30, no change if deprived at p60 (144); no change in noradrenergic fibre inputs nor beta-adrenergic receptors binding after in the SC after 1 month deprivation in p365 animals (145)<br>• <u>Activity:</u> reduced cytochrome oxidase activity in LGN and SC (90,146); 8 weeks after deprivation vLGN cytochrome oxidase activity recovered to control level while activity levels in LGN and SC remained lower, and 12 weeks after deprivation only SC had a significantly lower cytochrome oxidase activity (90); reduction in glucose uptake in SC and LGN up until 20 days after deprivation (55); increased binding of kainate, reduced binding of AMPA after 20 days (63); increase in choline acetyltransferase activity (SC;(92))<br>• <u>Protein expression:</u> increased substance-P immunoreactivity (LGN;(147)); increased expression of GAP-43 after 1 week and back to control levels after 2-3 weeks (SC;(148)); increased number thymosin beta 4- positive neurons and with longer and more complexly branched neurites (SC;(149))<br>2. **Principal neurons:** increase in the proportion of multi-innervated neurons and larger AMPAR-mediated EPSCs (dLGN;(71)); decrease in single-fibre AMPAR-mediated EPSC amplitudes at retinogeniculate synapse (LGN;(70,71) (Hooks & Chen, 2008b; Narushima et al., 2016)); no changes in the receptive fields in neurons of superficial SC cells (150)<br>3. **Interneurons:** fewer parvalbumin neurons in the LGN (101) and more parvalbumin neurons in the superficial SC (134); decrease in calbindin expression in vLGN (101) but no change in SC (134)<br>4. **Glia:** higher GFAP staining after 3 weeks with thick and branched processes visible (LGN;(122)); decrease in number of GFAP immunoreactive astrocytes in LGN after 12-48 weeks (101) |

## References

1. Boesveldt S, Postma EM, Boak D, Welge-Luessen A, Schöpf V, Mainland JD, et al. Anosmia—A Clinical Review. Chem Senses. 2017 Sep;42(7):513–23.

2. Worrall G. Acute otitis media. Can Fam Physician. 2007 Dec;53(12):2147–8.

3. Howarth A, Shone GR. Ageing and the auditory system. Postgrad Med J. 2006 Mar;82(965):166–71.

4. Ryan AF, Kujawa SG, Hammill T, Le Prell C, Kil J. Temporary and Permanent Noise-Induced Threshold Shifts: A Review of Basic and Clinical Observations. Otol Neurotol. 2016 Sep;37(8):e271–5.

5. Mullol J, Alobid I, Mariño-Sánchez F, Izquierdo-Domínguez A, Marin C, Klimek L, et al. The Loss of Smell and Taste in the COVID-19 Outbreak: a Tale of Many Countries. Curr Allergy Asthma Rep. 2020 Aug 3;20(10):61.

6. Muth CC. Sudden Vision Loss. JAMA. 2017 Aug 8;318(6):584.

7. Lepousez G, Valley MT, Lledo PM. The Impact of Adult Neurogenesis on Olfactory Bulb Circuits and Computations. Annual Review of Physiology. 2013;75(1):339–63.

8. Schwob JE, Jang W, Holbrook EH, Lin B, Herrick DB, Peterson JN, et al. Stem and progenitor cells of the mammalian olfactory epithelium: Taking poietic license. Journal of Comparative Neurology. 2017;525(4):1034–54.

9. Carulli D, Foscarin S, Rossi F. Activity-Dependent Plasticity and Gene Expression Modifications in the Adult CNS. Frontiers in Molecular Neuroscience [Internet]. 2011 [cited 2023 Aug 30];4.

10. Keck T, Toyoizumi T, Chen L, Doiron B, Feldman DE, Fox K, et al. Integrating Hebbian and homeostatic plasticity: the current state of the field and future research directions. Philos Trans R Soc Lond, B, Biol Sci. 2017 05;372(1715).

11. Turrigiano G. Too many cooks? Intrinsic and synaptic homeostatic mechanisms in cortical circuit refinement. Annu Rev Neurosci. 2011;34:89–103.

12. Wefelmeyer W, Puhl CJ, Burrone J. Homeostatic Plasticity of Subcellular Neuronal Structures: From Inputs to Outputs. Trends Neurosci. 2016;39(10):656–67.

13. Sancho L, Contreras M, Allen NJ. Glia as sculptors of synaptic plasticity. Neurosci Res. 2021 Jun;167:17–29.

14. Allen NJ, Lyons DA. Glia as architects of central nervous system formation and function. Science. 2018 Oct 12;362(6411):181–5.

15. Syka J. Plastic changes in the central auditory system after hearing loss, restoration of function, and during learning. Physiol Rev. 2002 Jul;82(3):601–36.

16. Chen JL, Lin WC, Cha JW, So PT, Kubota Y, Nedivi E. Structural basis for the role of inhibition in facilitating adult brain plasticity. Nat Neurosci. 2011 May;14(5):587–94.

17. Hubel DH, Wiesel TN. Receptive fields of cells in striate cortex of very young, visually inexperienced kittens. Journal of Neurophysiology. 1963 Nov;26(6):994–1002.

18. Feldman DE. Synaptic mechanisms for plasticity in neocortex. Annu Rev Neurosci. 2009;32:33–55.

19. Hensch TK. Critical period plasticity in local cortical circuits. Nat Rev Neurosci. 2005 Nov;6(11):877–88.

20. Duménieu M, Marquèze-Pouey B, Russier M, Debanne D. Mechanisms of Plasticity in Subcortical Visual Areas. Cells. 2021 Nov 13;10(11):3162.

21. Jones EG. Cortical and Subcortical Contributions to Activity-Dependent Plasticity in Primate Somatosensory Cortex. Annual Review of Neuroscience. 2000;23(1):1–37.

22. Maffei A, Turrigiano GG. Multiple modes of network homeostasis in visual cortical layer 2/3. J Neurosci. 2008 Apr 23;28(17):4377–84.

23. Pozo K, Goda Y. Unraveling mechanisms of homeostatic synaptic plasticity. Neuron. 2010 May 13;66(3):337–51.

24. Shepherd GM. The synaptic organization of the brain, 5th ed. New York, NY, US: Oxford University Press; 2004. xiv, 719 p. (The synaptic organization of the brain, 5th ed).

25. Kosaka T, Kosaka K. Neuronal organization of the main olfactory bulb revisited. Anat Sci Int. 2016 Mar;91(2):115–27.

26. Liu X, Huang H, Snutch TP, Cao P, Wang L, Wang F. The Superior Colliculus: Cell Types, Connectivity, and Behavior. Neurosci Bull. 2022 Dec;38(12):1519–40.

27. Weyand TG. The multifunctional lateral geniculate nucleus. Rev Neurosci. 2016 Feb;27(2):135–57.

28. Oertel D, Young ED. What’s a cerebellar circuit doing in the auditory system? Trends Neurosci. 2004 Feb;27(2):104–10.

29. Campagnola L, Manis PB. A map of functional synaptic connectivity in the mouse anteroventral cochlear nucleus. J Neurosci. 2014 Feb 5;34(6):2214–30.

30. Fox K, Stryker M. Integrating Hebbian and homeostatic plasticity: introduction. Philos Trans R Soc Lond, B, Biol Sci. 2017 05;372(1715).

31. Jaepel J, Hübener M, Bonhoeffer T, Rose T. Lateral geniculate neurons projecting to primary visual cortex show ocular dominance plasticity in adult mice. Nat Neurosci. 2017 Dec;20(12):1708–14.

32. Huh CYL, Abdelaal K, Salinas KJ, Gu D, Zeitoun J, Figueroa Velez DX, et al. Long-term Monocular Deprivation during Juvenile Critical Period Disrupts Binocular Integration in Mouse Visual Thalamus. J Neurosci. 2020 Jan 15;40(3):585–604.

33. Galliano E, Hahn C, Browne LP, R Villamayor P, Tufo C, Crespo A, et al. Brief Sensory Deprivation Triggers Cell Type-Specific Structural and Functional Plasticity in Olfactory Bulb Neurons. J Neurosci. 2021 Mar 10;41(10):2135–51.

34. Wilson D, Sullivan R. The D2 antagonist spiperone mimics the effects of olfactory deprivation on mitral/tufted cell odor response patterns. J Neurosci. 1995 Aug 1;15(8):5574–81.

35. Henegar JR, Maruniak JA. Quantification of the effects of long-term unilateral naris closure on the olfactory bulbs of adult mice. Brain Res. 1991 Dec 24;568(1–2):230–4.

36. Asako M, Holt AG, Griffith RD, Buras ED, Altschuler RA. Deafness-related decreases in glycine-immunoreactive labeling in the rat cochlear nucleus. J Neurosci Res. 2005 Jul 1;81(1):102–9.

37. Caicedo A, d’Aldin C, Eybalin M, Puel JL. Temporary sensory deprivation changes calcium-binding proteins levels in the auditory brainstem. J Comp Neurol. 1997 Feb 3;378(1):1–15.

38. Clarkson C, Antunes FM, Rubio ME. Conductive Hearing Loss Has Long-Lasting Structural and Molecular Effects on Presynaptic and Postsynaptic Structures of Auditory Nerve Synapses in the Cochlear Nucleus. J Neurosci. 2016 Sep 28;36(39):10214–27.

39. Francis HW, Manis PB. Effects of deafferentation on the electrophysiology of ventral cochlear nucleus neurons. Hear Res. 2000 Nov;149(1–2):91–105.

40. Garcia MM, Edward R, Brennan GB, Harlan RE. Deafferentation-induced changes in protein kinase C expression in the rat cochlear nucleus. Hear Res. 2000 Sep;147(1–2):113–24.

41. Poveda CM, Valero ML, Pernia M, Alvarado JC, Ryugo DK, Merchan MA, et al. Expression and Localization of Kv1.1 and Kv3.1b Potassium Channels in the Cochlear Nucleus and Inferior Colliculus after Long-Term Auditory Deafferentation. Brain Sci. 2020 Jan 8;10(1):35.

42. Whiting B, Moiseff A, Rubio ME. Cochlear nucleus neurons redistribute synaptic AMPA and glycine receptors in response to monaural conductive hearing loss. Neuroscience. 2009 Nov 10;163(4):1264–76.

43. Barmack NH, Qian Z. Activity-dependent expression of calbindin in rabbit floccular Purkinje cells modulated by optokinetic stimulation. Neuroscience. 2002;113(1):235–50.

44. Gonzalez-Perez O, López-Virgen V, Ibarra-Castaneda N. Permanent Whisker Removal Reduces the Density of c-Fos+ Cells and the Expression of Calbindin Protein, Disrupts Hippocampal Neurogenesis and Affects Spatial-Memory-Related Tasks. Front Cell Neurosci. 2018;12:132.

45. Philpot BD, Lim JH, Brunjes PC. Activity-dependent regulation of calcium-binding proteins in the developing rat olfactory bulb. J Comp Neurol. 1997 Oct 13;387(1):12–26.

46. Baker H, Kawano T, Margolis FL, Joh TH. Transneuronal regulation of tyrosine hydroxylase expression in olfactory bulb of mouse and rat. J Neurosci. 1983 Jan;3(1):69–78.

47. Andrade C. Understanding the Basics of Meta-Analysis and How to Read a Forest Plot: As Simple as It Gets. J Clin Psychiatry. 2020 Oct 6;81(5):20f13698.

48. Syka J, Rybalko N. Threshold shifts and enhancement of cortical evoked responses after noise exposure in rats. Hear Res. 2000 Jan;139(1–2):59–68.

49. Fokkens WJ, Bachert C, Douglas R, Gevaert P, Georgalas C, Harvey R, et al. European Position Paper on Rhinosinusitis and Nasal Polyps 2012. Rhinology. 2012;50(1–12):329.

50. Pula JH, Kwan K, Yuen CA, Kattah JC. Update on the evaluation of transient vision loss. Clin Ophthalmol. 2016;10:297–303.

51. Santos REA, da Silva MG, do Monte Silva MCB, Barbosa DAM, Gomes AL do V, Galindo LCM, et al. Onset and duration of symptoms of loss of smell/taste in patients with COVID-19: A systematic review. Am J Otolaryngol. 2021;42(2):102889.

52. Gold JR, Bajo VM. Insult-induced adaptive plasticity of the auditory system. Front Neurosci. 2014;8:110.

53. Park JW, Wang X, Xu RH. Revealing the mystery of persistent smell loss in Long COVID patients. Int J Biol Sci. 2022;18(12):4795–808.

54. Stitt AW, Curtis TM, Chen M, Medina RJ, McKay GJ, Jenkins A, et al. The progress in understanding and treatment of diabetic retinopathy. Prog Retin Eye Res. 2016 Mar;51:156–86.

55. Horsburgh K, McCulloch J. Alterations of functional glucose use and ligand binding to second messenger systems following unilateral orbital enucleation. Brain Res. 1991 May 24;549(2):317–21.

56. Wang WF, Kiyosawa M, Ishiwata K, Mochizuki M. Glucose metabolism in the visual structures of rat monocularly deprived by eyelid suture after postnatal eye opening. Jpn J Ophthalmol. 2005;49(1):6–11.

57. Briñón JG, Crespo C, Weruaga E, Martínez-Guijarro FJ, Aijón J, Alonso JR. Bilateral olfactory deprivation reveals a selective noradrenergic regulatory input to the olfactory bulb. Neuroscience. 2001;102(1):1–10.

58. Luo L, Ryan AF, Saint Marie RL. Cochlear ablation alters acoustically induced c-fos mRNA expression in the adult rat auditory brainstem. J Comp Neurol. 1999 Feb 8;404(2):271–83.

59. Barnes SJ, Sammons RP, Jacobsen RI, Mackie J, Keller GB, Keck T. Subnetwork-Specific Homeostatic Plasticity in Mouse Visual Cortex In Vivo. Neuron. 2015 Jun;86(5):1290–303.

60. Gainey MA, Feldman DE. Multiple shared mechanisms for homeostatic plasticity in rodent somatosensory and visual cortex. Philos Trans R Soc Lond, B, Biol Sci. 2017 05;372(1715).

61. Yan XX, Ribak CE. Increased expression of GABA transporters, GAT-1 and GAT-3, in the deafferented superior colliculus of the rat. Brain Res. 1998 Feb 2;783(1):63–76.

62. Li L, Gainey MA, Goldbeck JE, Feldman DE. Rapid homeostasis by disinhibition during whisker map plasticity. Proc Natl Acad Sci U S A. 2014 Jan 28;111(4):1616–21.

63. Chalmers DT, McCulloch J. Selective alterations in glutamate receptor subtypes after unilateral orbital enucleation. Brain Res. 1991 Feb 1;540(1–2):255–65.

64. Harris JA, Iguchi F, Seidl AH, Lurie DI, Rubel EW. Afferent deprivation elicits a transcriptional response associated with neuronal survival after a critical period in the mouse cochlear nucleus. J Neurosci. 2008 Oct 22;28(43):10990–1002.

65. Janz P, Illing RB. A role for microglial cells in reshaping neuronal circuitry of the adult rat auditory brainstem after its sensory deafferentation. J Neurosci Res. 2014 Apr;92(4):432–45.

66. Illing RB, Buschky H, Tadic A. Mitotic activity, modulation of DNA processing, and purinergic signalling in the adult rat auditory brainstem following sensory deafferentation. Eur J Neurosci. 2019 Dec;50(12):3985–4003.

67. Lau CG, Murthy VN. Activity-dependent regulation of inhibition via GAD67. J Neurosci. 2012 Jun 20;32(25):8521–31.

68. Tyler WJ, Petzold GC, Pal SK, Murthy VN. Experience-dependent modification of primary sensory synapses in the mammalian olfactory bulb. J Neurosci. 2007 Aug 29;27(35):9427–38.

69. Denizet M, Cotter L, Lledo PM, Lazarini F. Sensory deprivation increases phagocytosis of adult-born neurons by activated microglia in the olfactory bulb. Brain Behav Immun. 2017 Feb;60:38–43.

70. Hooks BM, Chen C. Vision triggers an experience-dependent sensitive period at the retinogeniculate synapse. J Neurosci. 2008 Apr 30;28(18):4807–17.

71. Narushima M, Uchigashima M, Yagasaki Y, Harada T, Nagumo Y, Uesaka N, et al. The Metabotropic Glutamate Receptor Subtype 1 Mediates Experience-Dependent Maintenance of Mature Synaptic Connectivity in the Visual Thalamus. Neuron. 2016 Sep 7;91(5):1097–109.

72. Louros SR, Hooks BM, Litvina L, Carvalho AL, Chen C. A role for stargazin in experience-dependent plasticity. Cell Rep. 2014 Jun 12;7(5):1614–25.

73. Hildebrandt H, Hoffmann NA, Illing RB. Synaptic reorganization in the adult rat’s ventral cochlear nucleus following its total sensory deafferentation. PLoS One. 2011;6(8):e23686.

74. Heusinger J, Hildebrandt H, Illing RB. Sensory deafferentation modulates and redistributes neurocan in the rat auditory brainstem. Brain Behav. 2019 Aug;9(8):e01353.

75. Grier BD, Belluscio L, Cheetham CEJ. Olfactory Sensory Activity Modulates Microglial-Neuronal Interactions during Dopaminergic Cell Loss in the Olfactory Bulb. Front Cell Neurosci [Internet]. 2016 Jul 15 [cited 2020 Mar 16];10. Available from: https://www.ncbi.nlm.nih.gov/pmc/articles/PM

76. Veyrac A, Didier A, Colpaert F, Jourdan F, Marien M. Activation of noradrenergic transmission by alpha2-adrenoceptor antagonists counteracts deafferentation-induced neuronal death and cell proliferation in the adult mouse olfactory bulb. Exp Neurol. 2005 Aug;194(2):444–56.

77. Gonzalez D, Satriotomo I, Miki T, Lee KY, Yokoyama T, Touge T, et al. Effects of monocular enucleation on calbindin-D 28k and c-Fos expression in the lateral geniculate nucleus in rats. Okajimas Folia Anat Jpn. 2005 May;82(1):9–18.

78. Förster CR, Illing RB. Plasticity of the auditory brainstem: cochleotomy-induced changes of calbindin-D28k expression in the rat. J Comp Neurol. 2000 Jan 10;416(2):173–87.

79. Fredrich M, Zeber AC, Hildebrandt H, Illing RB. Differential molecular profiles of astrocytes in degeneration and re-innervation after sensory deafferentation of the adult rat cochlear nucleus. Eur J Neurosci. 2013 Jul;38(1):2041–56.

80. Asher RA, Morgenstern DA, Fidler PS, Adcock KH, Oohira A, Braistead JE, et al. Neurocan is upregulated in injured brain and in cytokine-treated astrocytes. J Neurosci. 2000 Apr 1;20(7):2427–38.

81. Rowlands D, Lensjø KK, Dinh T, Yang S, Andrews MR, Hafting T, et al. Aggrecan Directs Extracellular Matrix-Mediated Neuronal Plasticity. J Neurosci. 2018 Nov 21;38(47):10102–13.

82. Huntley GW. Synaptic circuit remodelling by matrix metalloproteinases in health and disease. Nat Rev Neurosci. 2012 Nov;13(11):743–57.

83. Akol I, Kalogeraki E, Pielecka-Fortuna J, Fricke M, Löwel S. MMP2 and MMP9 Activity Is Crucial for Adult Visual Cortex Plasticity in Healthy and Stroke-Affected Mice. J Neurosci. 2022 Jan 5;42(1):16–32.

84. Hillen AEJ, Burbach JPH, Hol EM. Cell adhesion and matricellular support by astrocytes of the tripartite synapse. Prog Neurobiol. 2018;165–167:66–86.

85. Hamilton KA, Parrish-Aungst S, Margolis FL, Erdélyi F, Szabó G, Puche AC. Sensory deafferentation transsynaptically alters neuronal GluR1 expression in the external plexiform layer of the adult mouse main olfactory bulb. Chem Senses. 2008 Feb;33(2):201–10.

86. Maruniak JA, Taylor JA, Henegar JR, Williams MB. Unilateral naris closure in adult mice: atrophy of the deprived-side olfactory bulbs. Brain Res Dev Brain Res. 1989 May 1;47(1):27–33.

87. Pothayee N, Cummings DM, Schoenfeld TJ, Dodd S, Cameron HA, Belluscio L, et al. Magnetic resonance imaging of odorant activity-dependent migration of neural precursor cells and olfactory bulb growth. Neuroimage. 2017 Sep;158:232–41.

88. Cummings DM, Belluscio L. Continuous neural plasticity in the olfactory intrabulbar circuitry. J Neurosci. 2010 Jul 7;30(27):9172–80.

89. Fiske BK, Brunjes PC. Cell death in the developing and sensory-deprived rat olfactory bulb. J Comp Neurol. 2001 Mar 12;431(3):311–9.

90. Sukekawa K. Changes of cytochrome oxidase activity in the rat subcortical visual centers after unilateral eye enucleation. Neurosci Lett. 1987 Mar 31;75(2):127–32.

91. García del Caño G, Gerrikagoitia I, Martínez-Millán L. Plastic reaction of the rat visual cortico-collicular connection after contralateral retinal deafferentiation at the neonatal or adult stage: axonal growth versus reactive synaptogenesis. J Comp Neurol. 2002 Apr 29;446(2):166–78.

92. Gerrikagoitia I, García del Caño G, Martínez-Millán L. Quantifying presynaptic terminals at the light microscope level in intact and deafferented central nervous structures. Brain Res Brain Res Protoc. 2002 Jun;9(3):165–72.

93. Yuan Y, Shi F, Yin Y, Tong M, Lang H, Polley DB, et al. Ouabain-induced cochlear nerve degeneration: synaptic loss and plasticity in a mouse model of auditory neuropathy. J Assoc Res Otolaryngol. 2014 Feb;15(1):31–43.

94. Kawabata K, Maeda S, Takanaga A, Ito H, Tanaka K, Hayakawa T, et al. Apoptosis and retinal projections in the dorsal lateral geniculate nucleus after monocular deprivation during the later phase of the critical period in the rat. Anat Sci Int. 2003 Jun;78(2):104–10.

95. Carleton A, Petreanu LT, Lansford R, Alvarez-Buylla A, Lledo PM. Becoming a new neuron in the adult olfactory bulb. Nat Neurosci. 2003 May;6(5):507–18.

96. Bonzano S, Bovetti S, Gendusa C, Peretto P, De Marchis S. Adult Born Olfactory Bulb Dopaminergic Interneurons: Molecular Determinants and Experience-Dependent Plasticity. Front Neurosci. 2016;10:189.

97. Sawada M, Kaneko N, Inada H, Wake H, Kato Y, Yanagawa Y, et al. Sensory input regulates spatial and subtype-specific patterns of neuronal turnover in the adult olfactory bulb. J Neurosci. 2011 Aug 10;31(32):11587–96.

98. Bovetti S, Veyrac A, Peretto P, Fasolo A, De Marchis S. Olfactory enrichment influences adult neurogenesis modulating GAD67 and plasticity-related molecules expression in newborn cells of the olfactory bulb. PLoS One. 2009 Jul 23;4(7):e6359.

99. Yamaguchi M, Mori K. Critical period for sensory experience-dependent survival of newly generated granule cells in the adult mouse olfactory bulb. Proc Natl Acad Sci U S A. 2005 Jul 5;102(27):9697–702.

100. George NM, Gentile Polese A, Merle L, Macklin WB, Restrepo D. Excitable Axonal Domains Adapt to Sensory Deprivation in the Olfactory System. J Neurosci. 2022 Feb 23;42(8):1491–509.

101. Gonzalez D, Satriotomo I, Miki T, Lee KY, Yokoyama T, Touge T, et al. Changes of parvalbumin immunoreactive neurons and GFAP immunoreactive astrocytes in the rat lateral geniculate nucleus following monocular enucleation. Neurosci Lett. 2006 Mar 6;395(2):149–54.

102. Leist M, Datunashvilli M, Kanyshkova T, Zobeiri M, Aissaoui A, Cerina M, et al. Two types of interneurons in the mouse lateral geniculate nucleus are characterized by different h-current density. Sci Rep. 2016 Apr 28;6:24904.

103. Coppola DM. Studies of olfactory system neural plasticity: the contribution of the unilateral naris occlusion technique. Neural Plast. 2012;2012:351752.

104. Gudden. Experimentaluntersuchungen über das peripherische und centrale Nervensystem. Archiv f Psychiatrie. 1870 Oct 1;2(3):693–723.

105. Cheetham CEJ, Park U, Belluscio L. Rapid and continuous activity-dependent plasticity of olfactory sensory input. Nat Commun. 2016 Feb 22;7(1):1–11.

106. Bergman U, Ostergren A, Gustafson AL, Brittebo B. Differential effects of olfactory toxicants on olfactory regeneration. Arch Toxicol. 2002 Mar;76(2):104–12.

107. Cummings DM, Belluscio L. Charting plasticity in the regenerating maps of the mammalian olfactory bulb. Neuroscientist. 2008 Jun;14(3):251–63.

108. Hengen KB, Lambo ME, Van Hooser SD, Katz DB, Turrigiano GG. Firing rate homeostasis in visual cortex of freely behaving rodents. Neuron. 2013 Oct 16;80(2):335–42.

109. Kuhlman SJ, Olivas ND, Tring E, Ikrar T, Xu X, Trachtenberg JT. A disinhibitory microcircuit initiates critical-period plasticity in the visual cortex. Nature. 2013 Sep;501(7468):543–6.

110. Keck T, Scheuss V, Jacobsen RI, Wierenga CJ, Eysel UT, Bonhoeffer T, et al. Loss of sensory input causes rapid structural changes of inhibitory neurons in adult mouse visual cortex. Neuron. 2011 Sep 8;71(5):869–82.

111. Chambers AR, Resnik J, Yuan Y, Whitton JP, Edge AS, Liberman MC, et al. Central Gain Restores Auditory Processing following Near-Complete Cochlear Denervation. Neuron. 2016 Feb 17;89(4):867–79.

112. Henton A, Zhao Y, Tzounopoulos T. A Role for KCNQ Channels on Cell Type-Specific Plasticity in Mouse Auditory Cortex after Peripheral Damage. J Neurosci. 2023 Mar 29;43(13):2277–90.

113. Kumar M, Handy G, Kouvaros S, Zhao Y, Brinson LL, Wei E, et al. Cell-type-specific plasticity of inhibitory interneurons in the rehabilitation of auditory cortex after peripheral damage. Nat Commun. 2023 Jul 13;14(1):4170.

114. Gainey MA, Aman JW, Feldman DE. Rapid Disinhibition by Adjustment of PV Intrinsic Excitability during Whisker Map Plasticity in Mouse S1. J Neurosci. 2018 May 16;38(20):4749–61.

115. Jamann N, Dannehl D, Lehmann N, Wagener R, Thielemann C, Schultz C, et al. Sensory input drives rapid homeostatic scaling of the axon initial segment in mouse barrel cortex. Nat Commun. 2021 Jan 4;12:23.

116. Dan Y, Poo MM. Spike timing-dependent plasticity of neural circuits. Neuron. 2004 Sep 30;44(1):23–30.

117. Gambino F, Holtmaat A. Spike-timing-dependent potentiation of sensory surround in the somatosensory cortex is facilitated by deprivation-mediated disinhibition. Neuron. 2012 Aug 9;75(3):490–502.

118. Karmarkar UR, Buonomano DV. Different forms of homeostatic plasticity are engaged with distinct temporal profiles. Eur J Neurosci. 2006 Mar;23(6):1575–84.

119. Kullmann DM, Moreau AW, Bakiri Y, Nicholson E. Plasticity of inhibition. Neuron. 2012 Sep 20;75(6):951–62.

120. Byrne DJ, Lipovsek M, Crespo A, Grubb MS. Brief sensory deprivation triggers plasticity of dopamine-synthesising enzyme expression in genetically labelled olfactory bulb dopaminergic neurons. European Journal of Neuroscience. 2022;56(1):3591–612.

121. Linden ML, Heynen AJ, Haslinger RH, Bear MF. Thalamic activity that drives visual cortical plasticity. Nat Neurosci. 2009 Apr;12(4):390–2.

122. Canady KS, Olavarria JF, Rubel EW. Reduced retinal activity increases GFAP immunoreactivity in rat lateral geniculate nucleus. Brain Res. 1994 Nov 14;663(2):206–14.

123. Kikuta S, Sakamoto T, Nagayama S, Kanaya K, Kinoshita M, Kondo K, et al. Sensory deprivation disrupts homeostatic regeneration of newly generated olfactory sensory neurons after injury in adult mice. J Neurosci. 2015 Feb 11;35(6):2657–73.

124. Mandairon N, Jourdan F, Didier A. Deprivation of sensory inputs to the olfactory bulb up-regulates cell death and proliferation in the subventricular zone of adult mice. Neuroscience. 2003;119(2):507–16.

125. Mast TG, Fadool DA. Mature and precursor brain-derived neurotrophic factor have individual roles in the mouse olfactory bulb. PLoS One. 2012;7(2):e31978.

126. Mostafapour SP, Del Puerto NM, Rubel EW. bcl-2 Overexpression eliminates deprivation-induced cell death of brainstem auditory neurons. J Neurosci. 2002 Jun 1;22(11):4670–4.

127. Jin YM, Godfrey DA, Sun Y. Effects of cochlear ablation on choline acetyltransferase activity in the rat cochlear nucleus and superior olive. J Neurosci Res. 2005 Jul 1;81(1):91–101.

128. Tucci DL, Cant NB, Durham D. Effects of conductive hearing loss on gerbil central auditory system activity in silence. Hear Res. 2001 May;155(1–2):124–32.

129. Illing RB, Horváth M. Re-emergence of GAP-43 in cochlear nucleus and superior olive following cochlear ablation in the rat. Neurosci Lett. 1995 Jul 14;194(1–2):9–12.

130. Zilles K, Wree A, Petrovic-Minic B, Schleicher A, Beck T. Different metabolic changes in the lateral geniculate nucleus and the superior colliculus of adult rats after simultaneous or delayed double enucleation. Brain Res. 1989 May 29;488(1–2):14–21.

131. Giraldi-Guimarães A, Mendez-Otero R. Induction of the candidate-plasticity NGFI-A protein in the adult rat superior colliculus after visual stimulation. Brain Res Mol Brain Res. 2005 Feb 18;133(2):242–52.

132. Vierci G, Oliveira CSD, Perera LR, Bornia N, Leal RB, Rossi FM. Creb is modulated in the mouse superior colliculus in developmental and experimentally-induced models of plasticity. Int J Dev Neurosci. 2013 Feb;31(1):46–52.

133. Boulenguez P, Abdelkefi J, Pinard R, Christolomme A, Segu L. Effects of retinal deafferentation on serotonin receptor types in the superficial grey layer of the superior colliculus of the rat. J Chem Neuroanat. 1993;6(3):167–75.

134. Lane RD, Allan DM, Bennett-Clarke CA, Rhoades RW. Differential age-dependent effects of retinal deafferentation upon calbindin- and parvalbumin-immunoreactive neurons in the superficial layers of the rat’s superior colliculus. Brain Res. 1996 Nov 18;740(1–2):208–14.

135. Gribaudo S, Bovetti S, Friard O, Denorme M, Oboti L, Fasolo A, et al. Transitory and activity-dependent expression of neurogranin in olfactory bulb tufted cells during mouse postnatal development. J Comp Neurol. 2012 Oct 1;520(14):3055–69.

136. Wilson DA, Wood JG. Functional consequences of unilateral olfactory deprivation: time-course and age sensitivity. Neuroscience. 1992 Jul;49(1):183–92.

137. Marks CA, Cheng K, Cummings DM, Belluscio L. Activity-dependent plasticity in the olfactory intrabulbar map. J Neurosci. 2006 Nov 1;26(44):11257–66.

138. Kosaka T, Katsumaru H, Hama K, Wu JY, Heizmann CW. GABAergic neurons containing the Ca2+-binding protein parvalbumin in the rat hippocampus and dentate gyrus. Brain Res. 1987 Sep 1;419(1–2):119–30.

139. Parrish-Aungst S, Kiyokage E, Szabo G, Yanagawa Y, Shipley MT, Puche AC. Sensory experience selectively regulates transmitter synthesis enzymes in interglomerular circuits. Brain Res. 2011 Mar 25;1382:70–6.

140. Pérez-Revuelta L, Téllez de Meneses PG, López M, Briñón JG, Weruaga E, Díaz D, et al. Secretagogin expression in the mouse olfactory bulb under sensory impairments. Sci Rep. 2020 Dec 9;10(1):21533.

141. Kelsch W, Lin CW, Mosley CP, Lois C. A critical period for activity-dependent synaptic development during olfactory bulb adult neurogenesis. J Neurosci. 2009 Sep 23;29(38):11852–8.

142. Heeringa AN, Stefanescu RA, Raphael Y, Shore SE. Altered vesicular glutamate transporter distributions in the mouse cochlear nucleus following cochlear insult. Neuroscience. 2016 Feb 19;315:114–24.

143. Rhoades RW, Mooney RD, Chiaia NL, Bennett-Clarke CA. Development and plasticity of the serotoninergic projection to the hamster’s superior colliculus. J Comp Neurol. 1990 Sep 8;299(2):151–66.

144. Nakamura S, Shirokawa T, Sakaguchi T. Increased projection from the locus coeruleus to the lateral geniculate nucleus and visual cortex in young adult rats following unilateral enucleation. Neurosci Lett. 1984 Aug 24;49(1–2):77–80.

145. Muguruma K, Matsumura K, Watanabe Y, Shiomitsu T, Imamura K, Watanabe Y. Effects of monocular enucleation on receptor binding and innervation pattern of the noradrenergic system in the superior colliculus of the pigmented rat. Neurosci Res. 1997 Aug;28(4):311–24.

146. Land PW. Dependence of cytochrome oxidase activity in the rat lateral geniculate nucleus on retinal innervation. J Comp Neurol. 1987 Aug 1;262(1):78–89.

147. Senba E, Miguell-Hidalgo JJ. Substance P in the retina and primary visual centers: its projection and plasticity after deafferentation. Regul Pept. 1993 Jul 2;46(1–2):129–37.

148. Mendonça HR, Araújo SES, Gomes ALT, Sholl-Franco A, da Cunha Faria Melibeu A, Serfaty CA, et al. Expression of GAP-43 during development and after monocular enucleation in the rat superior colliculus. Neurosci Lett. 2010 Jun 14;477(1):23–7.

149. Paulussen M, Arckens L. Striking neuronal thymosin beta 4 expression in the deep layers of the mouse superior colliculus after monocular deprivation. Brain Struct Funct. 2012 Jan;217(1):81–91.

150. Carrasco MM, Pallas SL. Early visual experience prevents but cannot reverse deprivation-induced loss of refinement in adult superior colliculus. Vis Neurosci. 2006;23(6):845–52.

